# BONCAT-Live for isolation and cultivation of active environmental microbes

**DOI:** 10.1101/2025.05.14.654084

**Authors:** Sayali A. Mulay, Tatiana A. Vishnivetskaya, Leah H. Hochanadel, Dawn M. Klingeman, Karen G. Lloyd, Dale A. Pelletier, Mircea Podar

## Abstract

In diverse environments, microbes drive a myriad of processes, from geochemical and nutrient cycling to interspecies interactions, including in association with plants and animals. Their physiological state is dynamic and impacted by abiotic and biotic conditions, responding to environmental fluctuations by changes in cellular metabolism, according to their genetic potential. Molecular, cellular and genomic approaches can identify and measure microbial responses and adaptation to environmental changes in native communities. However, isolating the individual microbes that respond to specific changes has been difficult. To address that, we implemented BONCAT-Live, by integrating bio-orthogonal non-canonical amino acid tagging (BONCAT) in diverse native communities, with isolation and cultivation of cells responding to specific stimuli, at different time scales. In frozen Arctic permafrost samples, we identified and isolated dormant bacteria that become active after thawing under native or nutrient enriched conditions. From a *Populus* tree rhizosphere, we isolated strains that thrive under high concentrations of root exudates that act as defense compounds and nutrients. In the human oral microbiome, we identified and isolated bacteria that rapidly proliferated when exposed to metabolites provided by the host or other co-occurring microbes. Further characterization of isolated bacterial strains will provide opportunities for in depth determination of how these microbes adapt to changes in their environments, individually and as part of model communities.

**Importance:** Dynamic microbial activity transforms environments and impacts health and disease in associations with plants and animals, including humans. Identifying the contribution of individual microbes to those processes in real time has not been generally compatible with their selective cultivation. BONCAT-Live tracks which microbes in environmental samples are translationally active and couples it with single cell isolation and cultivation. By studying the response of individual community member to specific natural or induced physical or chemical changes in the environment and culturing those organisms, BONCAT-Live enables further insights into microbial metabolic strategies, community dynamics and environmental adaptations.

## INTRODUCTION

Most environments harbor complex microbial communities, taxonomically diverse, partitioned across various niches and ranging from specialists to physiologically versatile. Fluctuations of physical, chemical and biotic parameters, at different spatial and temporal scales, can selectively impact and partition the metabolic states of individual members of the community. Taxa that are under permissive conditions (temperature, nutrients, energy sources etc.), would display various levels of physiological activity (e.g. respiration, protein synthesis, cell division), while those facing adversities and limitations may enter dormancy (including sporulation, viable but non-culturable state)(1–3) until optimality resumes. Physiological activation from dormant states (as evidenced by gene expression, protein synthesis, cell division, motility etc.), can occur rapidly, over minutes or hours, even after long periods of time (4, 5). In some microbes, metabolic activity can persist across a wide range of conditions (e.g. low temperature-adapted microbes) by adopting distinct strategies (genes, metabolites) at different states (6–8). At the community level, a wide range of techniques can detect and quantify specific biochemical processes (e.g. respiration, denitrification, methanogenesis, sulfate reduction etc.), in situ and in the laboratory. By using stable isotope probing coupled with metagenomic sequencing, physiologically active microbes can be identified and quantified (9) and, when combined with spectroscopic techniques, dissected at cellular and subcellular levels (10, 11). A variety of metabolic substrate analogs have been developed to track specific cellular processes and how microbes react under changing conditions (10). Cell surface labelling using click-chemistry has been demonstrated by using analogs incorporated into the peptidoglycan and the outer membrane (12–14). However, because of the many differences in the structure and cellular envelope synthesis across microbes, there is no single, broadly applicable substrate for environmental tagging. BONCAT uses amino acid analogs that can be conjugated via click reactions to chemical tags (e.g. fluorophores), to label proteins in translationally active cells (15, 16), including in environmental samples (17–22). Those cells can subsequently be identified and visualized via microscopy-fluorescence *in situ* hybridization (FISH) or isolated by flow cytometry cell sorting (BONCAT-FACS) followed by amplicon sequencing (20, 21). So far and to our knowledge BONCAT has only been applied as an end-point analysis, without recovering viable microbes for subsequent studies.

Microbial isolation and cultivation, whether by traditional or high throughput approaches, relies on using a broad range of conditions, from highly selective to broadly permissive in nutrients, energy sources, and temperature (e.g. (23–29)). That leads to isolates that may or may not have been highly active in the environment at the time of collection or indicative of their responses to changing conditions or interactions with other species. Here, we have implemented BONCAT-Live as an approach to isolate and culture translationally active bacteria that respond to simulated environmental changes at different time scales, from weeks to hours. Specifically, we modeled permafrost thawing, the responses of *Populus* tree rhizosphere to plant-secreted metabolites and that of human oral microbiota to host and community shared nutrients.

## RESULTS

### Cell surface click-chemistry and viability tests enable BONCAT-Live

Previous studies that applied BONCAT to characterize active microbial populations have primarily focused on labeling and detection efficiency. Because the reagents used in the click-chemistry reactions may not effectively pass through the cellular membranes, fixation has been routinely used to enable access to cytoplasmic proteins, which are expected to be the major targets for tagging. Testing whether surface exposed proteins may be sufficient substrates for click-chemistry tagging of cells with an intact membrane was an important step in evaluating BONCAT-Live feasibility (**Figure 1**). We grew pure cultures of six phylogenetically diverse *Populus* associated soil bacteria (*Terriglobus* sp., *Flavobacterium* sp., *Bacillus* sp., *Rhizobium* sp., *Paraburkholderia* sp. and *Roseimicrobium* sp.)(30) in the presence of homopropargylglycine (HPG) and performed fixation-independent copper (Cu)-mediated click reactions with a biotin-azide tag. Microscopic examination after labeling with fluorescent streptavidin revealed successful tagging (**Figure 2**). To determine if the biotin tags are accessible on the cellular surface, we seeded those six HPG-containing bacteria to a live *Populus* rhizosphere soil microbiota sample and, after click reaction with biotin-azide, we used streptavidin-coated magnetic beads to capture labeled cells. SSU rRNA gene V4 amplicon sequencing of the samples, before and after magnetic capture shows that the seeded bacteria were preferentially enriched from the highly diverse rhizosphere sample (**Figure 2**). While not intended to be a quantitative assay, this demonstrates the accessibility of HPG-containing surface proteins to tagging for detection and isolation.

**Figure 1.**
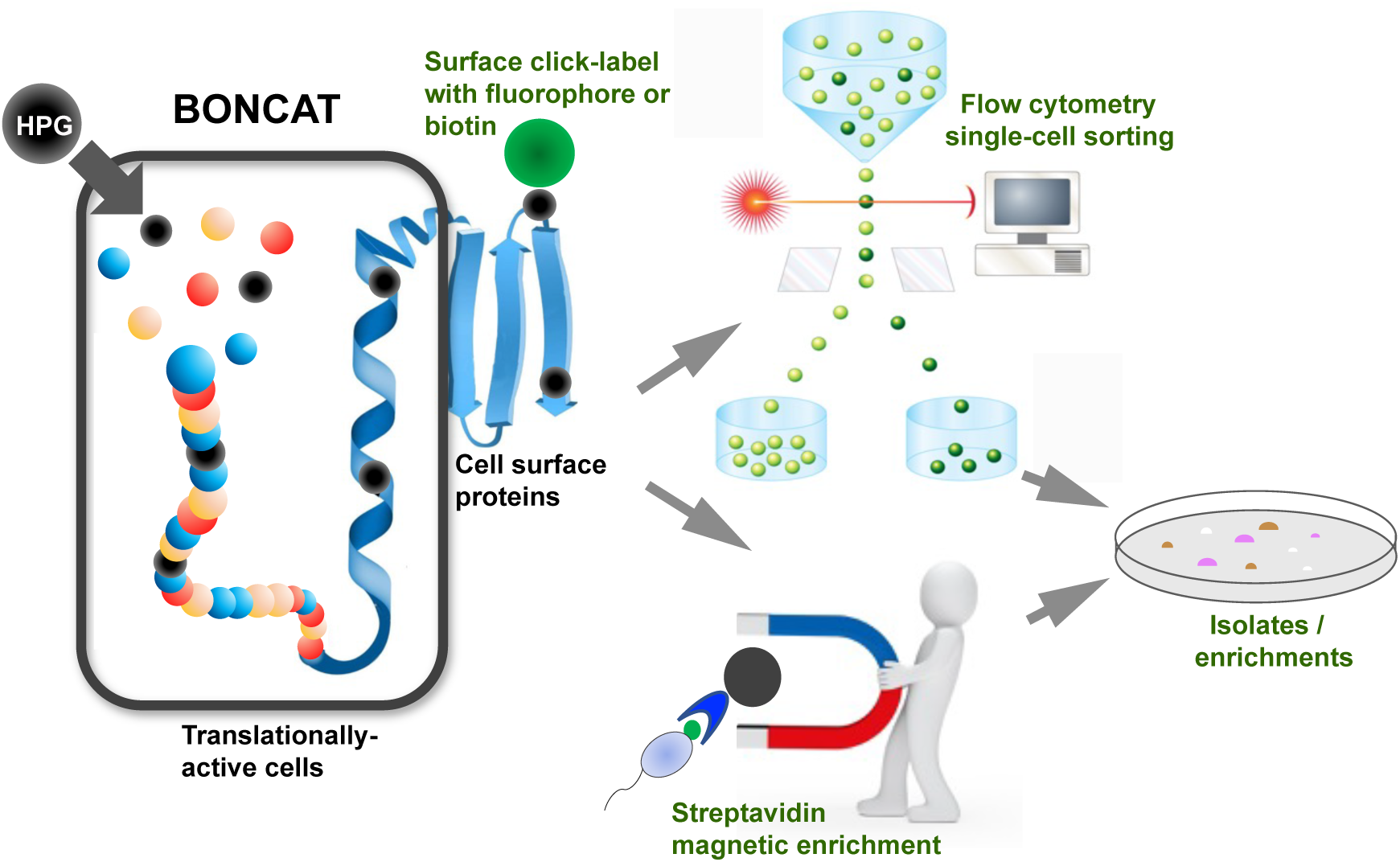
Concept and workflow diagram for BONCAT-Live.

**Figure 2.**
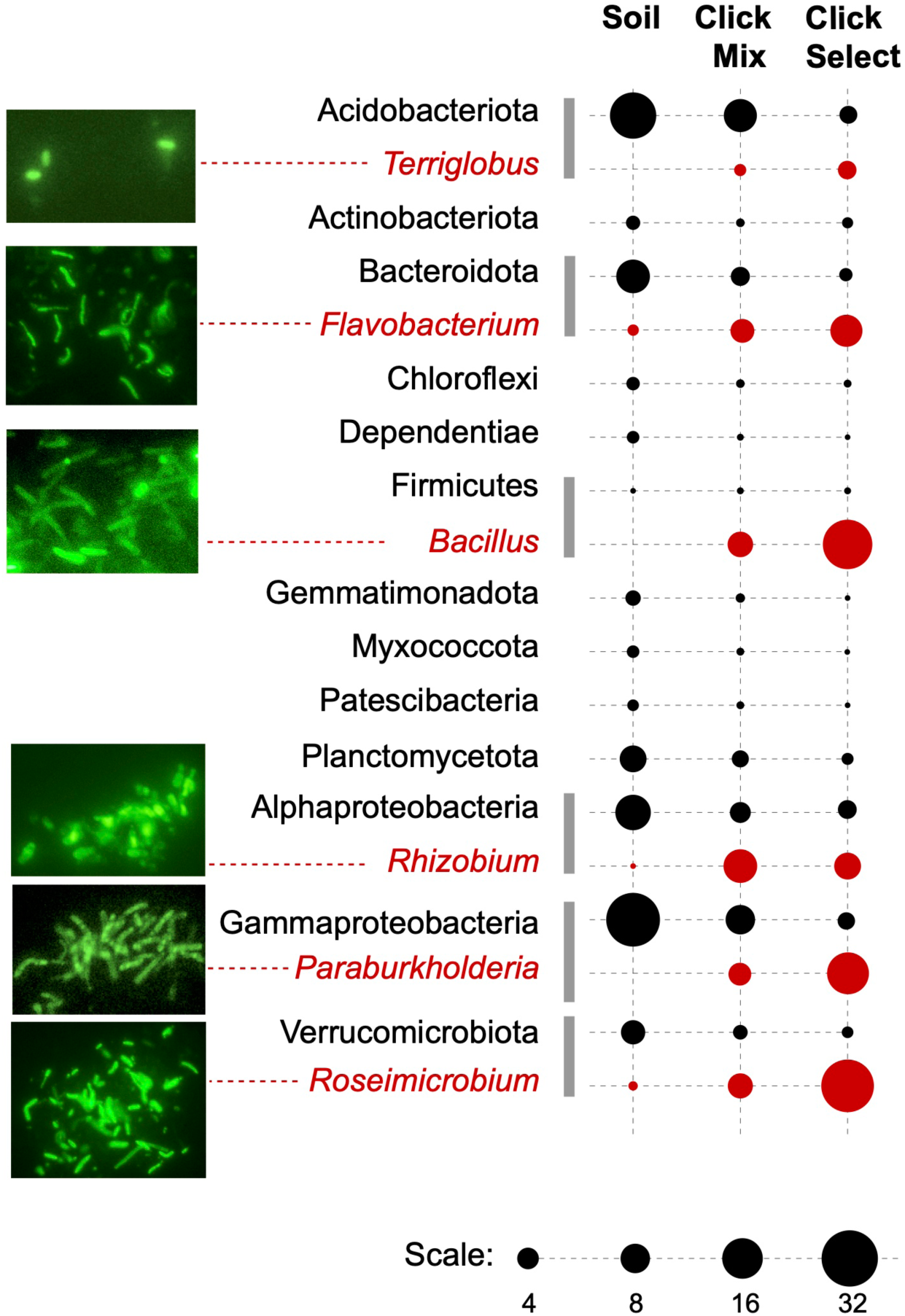
Cell surface BONCAT labeling and selection. Strains of six bacterial genera (red) were labeled with biotin after BONCAT and imaged after surface staining with Streptavidin-A488 or magnetically enriched after mixing in a soil microbiota sample. The bubble plot indicates taxon relative abundance based on SSU rRNA V4 amplicons comparing the soil sample before and after seeding with the magnetically enriched eluate.

Next, we investigated how the treatment steps and chemicals associated with BONCAT, click chemistry and flow cytometry cell sorting impact microbial viability, as each subjects the cells to different stresses. While we recognize that the many species of microbes present in the environment would likely respond differently, we evaluated those parameters primarily using a reference *Pseudomonas* sp. GM41 isolated from *Populus deltoides* rhizosphere (30). The assay end point was colony formation after flow sorting of 100 individual cells onto R2A nutrient agar (**Supplementary Figure S1**). Flow sorting itself did not have a major impact on viability, as we consistently obtained 80-90% viability, with or without including a DNA staining dye (Syto13 or Syto61). Similarly, the addition of either azidohomoalanine (AHA, 100μM) or HPG (50μM) during culture incubation did not noticeably affect viability (>80%, **Supplementary Figure S1**), supporting prior reports of low toxicity in pure cultures and environmental samples at those concentrations (15, 31, 32). Bio-orthogonal amino acids can have different toxic effects on some organisms, depending on concentration and culture conditions (33). We next investigated the effect of click labeling reactions on viability. Strain-promoted, Cu-independent chemistry using a dibenzocyclooctyne (DBCO)-containing dye requires blocking of free thiols with a high concentration of 2-chloroacetamide (>10 mM), which led to poor cell viability (<20%), so this process was not suitable for our test environments. We instead used Cu-mediated click-chemistry coupled with fluorescent AZDye 488 picolyl azide cellular labeling following incubations with HPG. The individual compounds necessary to generate the reduced environment and block non-specific reactions (ascorbate and aminoguanidine) were not significantly toxic at the standard concentrations (up to 5 mM) and incubation times (30-60 min) for click-labeling reactions. The typically used Cu concentration (50 μM) had a more detrimental impact on viability, with >50% inhibition (**Supplementary Figure S1**). While an essential microelement, Cu can be toxic by generating reactive oxygen species (ROS), inactivating iron-sulfur clusters in essential enzymes, and microbes have evolved various mechanisms to counteract it (34, 35). After tests to mitigate Cu toxicity during click reactions, we found that reducing its concentration to 5 μM and including ROS scavengers (pyruvate and catalase) maintained efficient click labeling while improving viability to ∼80%, conditions we further applied to the BONCAT-Live studies. Pyruvate and catalase have been previously found to improve microbial viability and cultivability (36–38) and may be especially beneficial for organisms that are sensitive to ROS.

### Microbes in the *Populus* rhizosphere are stimulated by root exudates

Plants establish symbiotic associations with microbes and fungi in the rhizosphere and within the roots by diverse chemical cues that reshape the diversity relative to the surrounding soil (39–42). Among the compounds that plant roots exude at various levels are amino acids, sugars, and organic acids which are used as nutrients by the surrounding microbiota. In addition, plant produce specific defense compounds to counteract pathogens and grazers. We used freshly collected *Populus deltoides* fine roots (2-3 mm thick) with the immediately surrounding soil. Root exudates often form a concentration gradient that is dependent on the amount and rate of secretion, soil diffusion and rate of utilization by different microorganisms. Some compounds, including antimicrobial secondary metabolites, can be utilized sequentially by distinct microbial guilds and provide nutritional handoffs (cross-feeding) to other taxa (42, 43). Also, under native environmental conditions, limited by water, nutrient availability and competition/predation, microbes have variable effective growth/division rates, ranging from hours to days (44). Based on preliminary tests we detected BONCAT-positive rhizosphere bacteria within 12-48 hours of incubation. Within the same time frame, prior studies have applied BONCAT to identify active microbes under native or stimulated conditions of various soils (20, 45).

The *Populus* rhizosphere samples presented a rich microbial diversity at all taxa levels (>30 different phyla and >500 genera/species) (**Figure 3, Supplemental Table**). Background control incubations (fixed sample or without HPG) resulted in no label being incorporated by click-chemistry, as determined by flow cytometry. Incubations in the presence of HPG for 24 hours but with no exogenous growth substrates, revealed a translationally active population of microbes (∼2% of the DNA-positive particles) relying on soil and root-derived nutrients. Based on rRNA amplicon sequencing of labeled and sorted cells, alpha- and gamma-proteobacteria dominated, with members of the *Sphingomonadaceae* having the highest relative abundance. Based on this background level of activity, we supplemented the incubations with malate, a root-produced organic acid known to promote recruitment of beneficial bacteria (46). Following 24 hours, ∼7.5% of the cells were BONCAT-positive, dominated by *Pseudomonas* strains, as determined by amplicon sequencing (**Figure 3**). From this population of translationally active cells, we flow-sorted single cells on R2A nutrient agar plates and incubated them to form colonies (**Figure 4**). Based on full length 16S rRNA gene amplification and sequencing, the isolates were assigned to *Pseudomonas*, other proteobacteria (*Pantoea, Ralstonia, Sphingomonas*), as well as several members of Actinobacteriota and Bacteroidota (**Figure 4**) known to associate with plant roots, including *Populus* (30).

**Figure 3.**
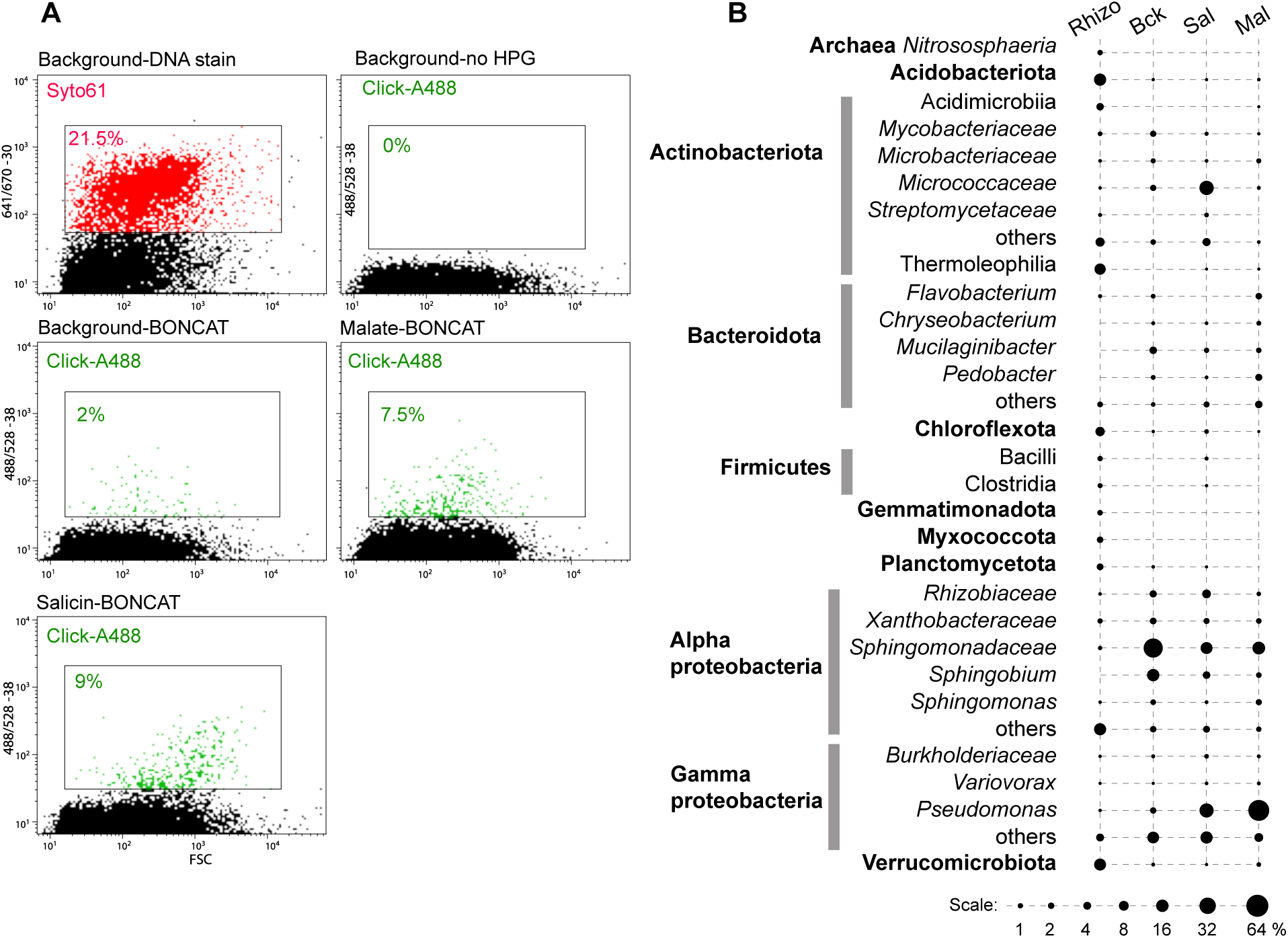
Rhizosphere BONCAT-Live experiment. **(A)** *Populus deltoides* rhizosphere samples were incubated for BONCAT with no exogenous nutrients (Background), with salicin (Sal) or malate (Mal), followed by click-labeling with Alexa488, DNA staining (Syto61) and flow cytometry cel sorting. Cytometry plots and color gates indicate frequency of labeled particles for control and BONCAT-labeled samples (DNA staining not shown for all samples). **(B)** Microbial taxa relative abundance in BONCAT-positive sorted cell populations based on gating shown in (A).

**Figure 4.**
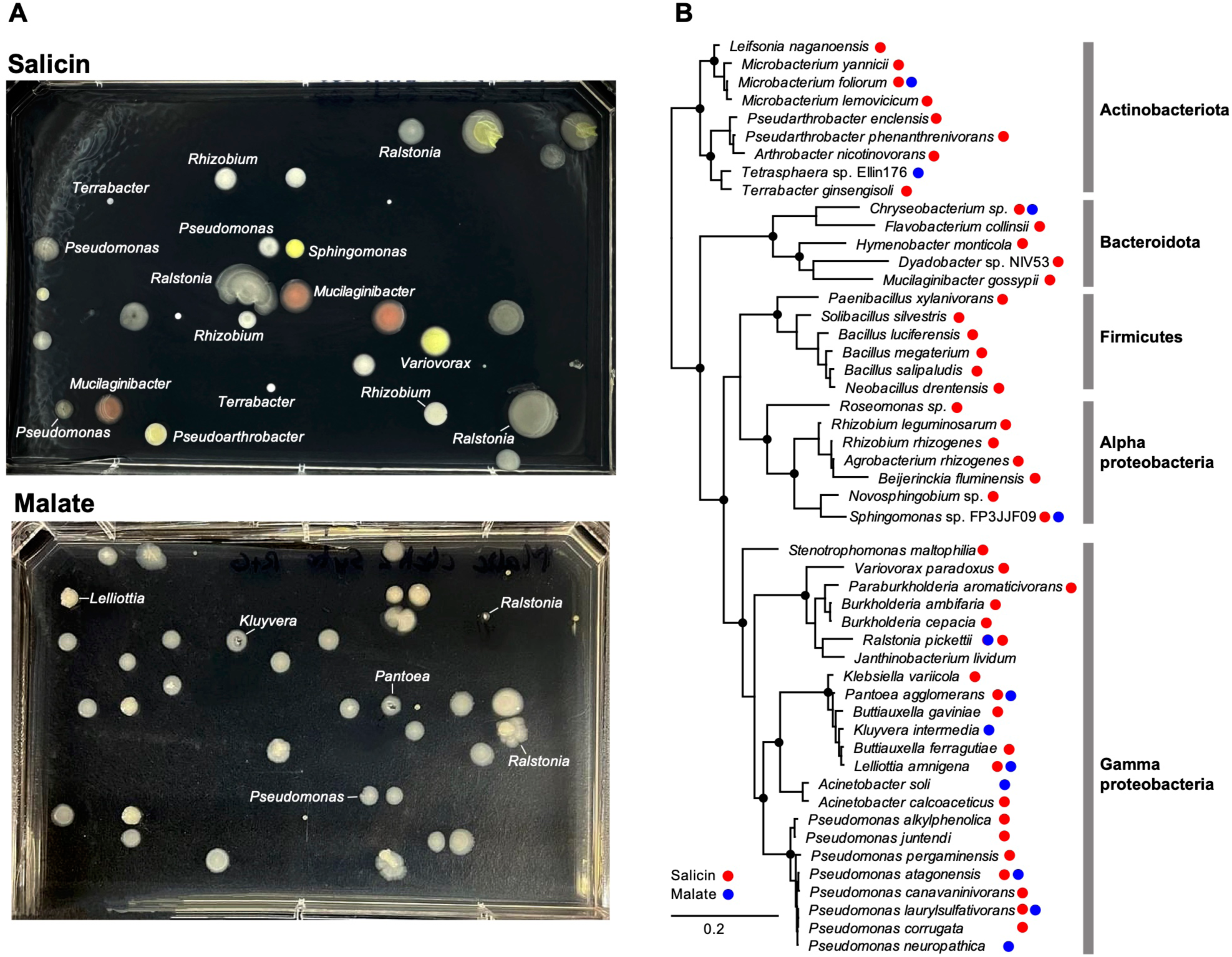
Microbial isolates from rhizosphere BONCAT-Live. **(A)** Single cells from salicin and malate gates were sorted on R2A agar and incubated to form colonies. Shown are example plates and identity of selected bacterial colonies based on SSU rRNA amplicon sequencing. **(B)** SSU rRNA phylogeny of bacterial species for which strains were isolated from salicin and malate BONCAT-Live.

From among the known *Populus* defense/regulatory exudates (47) we selected salicin, representing the salicylic acid (SA) plant hormonal and defense pathway (48, 49). Salicin is an abundant phenolic glycoside in *Populus* and through stepwise enzymatic breakdown and cross-feeding amongst different bacteria (e.g., *Rahnella*, *Pseudomonas*), it has been proposed to contribute to the root microbiota assembly (43). Incubation of the rhizosphere-root samples with salicin resulted in 9% BONCAT-positive cells (**Figure 3**). The highest increase in relative abundance, based on amplicon sequencing, was among Actinobacteriota (especially the *Micrococcaceae*), as well as *Pseudomonas* and other proteobacteria. Sorting of single cells on non-selective R2A agar media yielded two dozen distinct strains representing a rich diversity of Actinobacteriota, Bacteroidota, Firmicutes and proteobacteria, close to many known root symbionts (**Figure 4**).

### BONCAT-Live to study Arctic permafrost thawing

Permafrost is perennially frozen ground that covers a fifth of the Earth’s land surface (50) and stores half of the global soil organic carbon (51, 52). The permafrost lies beneath a surface soil layer that undergoes seasonal freeze-thaw cycles (active layer). As a result of global warming, the active layer that is deepening and thawing reaches into the permafrost. Microbes that have been frozen for up to millions of years can become metabolically active and breakdown organic matter, releasing greenhouse gases into the atmosphere and creating a positive feedback loop (53). As part of a broader study of permafrost communities, we analyzed cores collected from the Bayelva site at Ny Ålesund, Svalbard (Norway), location that is experiencing more extreme climate changes than the rest of the Arctic (54). In 2021, the recorded active layer temperature at this site was >0°C, with temperature maximum of ∼8°C (55). While a variety of biogeochemical and metagenomic approaches have been used to study thawing permafrost and its microbiota (56, 57), BONCAT has not been reported so far, to our knowledge. Additionally, BONCAT-Live could enable direct access to microbial strains as they become physiologically active under thawing conditions.

Samples from frozen cores that spanned the active layer (about 0.5-0.7 m depth) and the deep permafrost (>2 m depth) had distinct microbial composition. Some of the more typical soil taxa (including Acidobacteriota, Actinobacteriota, Gemmatimonadota, Chloroflexi, *Sphingomonas,* Verrucomicrobiota) were represented in the active layers, while the permafrost had higher levels of various proteobacteria (*Aliihoeflea, Enhydrobacter, Pseudomonas*) as well as Myxococcota and uncultured Actinobacteriota (**Figure 5**). We tested the effects of incubation temperature (−20°C, 4°C) and time (2-6 weeks at 4°C, 12 weeks at −20°C) on BONCAT output (fluorescent particles following click-labeling), cultivation temperature after cell sorting (15°C or 25°C) and media (standard/diluted nutrient agar). Because the site is characterized as a high lithic and low nutrient permafrost and permafrost-affected soils (58), we also supplemented some of the incubations with defined R2A-type nutrients.

**Figure 5.**
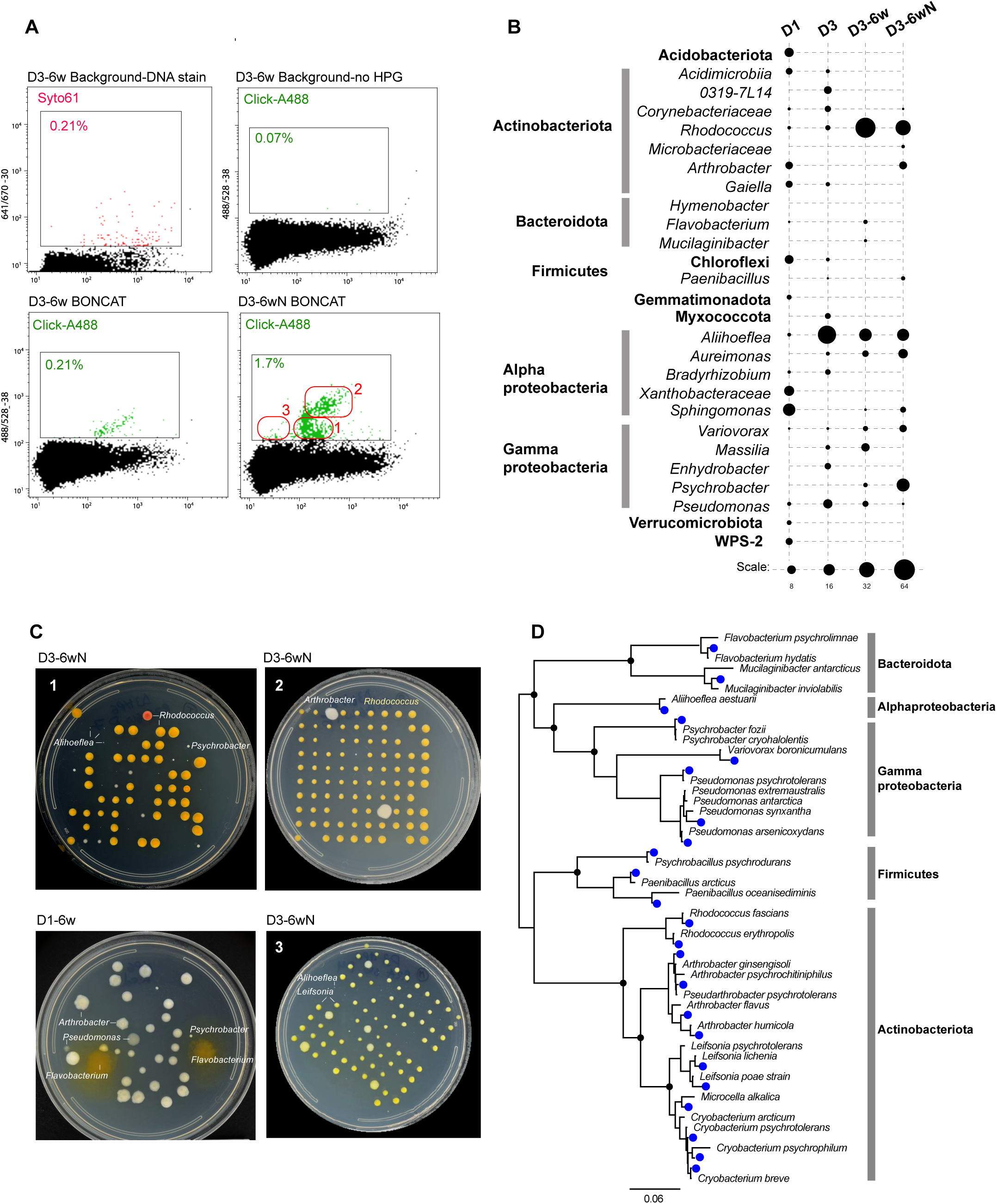
BONCAT-Live on Arctic permafrost cores. Samples from the active layer (D1) and permafrost (D3) were incubated for BONCAT without or with exogenous nutrients (N). **(A)** Flow cytometry plots for permafrost (D3) BONCAT, stained for DNA or click-labeled with Alexa488. Gates used for sorting and particle frequency are indicated. Nutrient-amended sample was fine gated (1, 2, 3) for bacterial isolation. **(B)** Microbial taxa relative abundance in original core samples (D1-D3) and BONCAT-positive sorted cell populations based on gating shown in (A). **(C)** Bacterial isolates following BONCAT-positive single cell sorting from non-amended and nutrient-enriched (N) permafrost incubations, based on gates shown in (A). Identity of representative types of colonies is indicated. **(D)** SSU rRNA phylogeny of isolated bacterial strains (blue dots) and their closest relative described species.

The incubations performed at −20°C, as well as the negative BONCAT-controls, yielded no click-labeling signal. The BONCAT-Live experiments were conducted mainly on 6-week incubations using permafrost layer samples and were performed twice, with subsamples from the same main core. During flow sorting analysis of the click-labeled samples, we sorted both entire labeled populations as well as sub-populations based on gates that suggested distinct scatter and fluorescence properties (**Figure 5**). Based on amplicon sequencing of BONCAT-positive populations, specific taxa increased significantly in abundance relative to the starting permafrost. They included *Rhodococcus* and *Arthrobacter* (Actinobacteriota), *Aliihoeflea* and *Aureimonas* (Alphaproteobacteria) and *Psychrobacter* (Gammaproteobacteria). Supplementing the incubations with exogenous R2A-type nutrients had a relatively minor effect on alpha diversity, although nutrients stimulated some taxa and inhibited others, which was also evidenced in flow cytometry profiles (**Figure 5**). Viability after single cell deposition on media plates was comparable to what we had determined with tests on soil bacteria and with a *Pseudomonas* strain previously isolated from the same site (57). High viability was reflected in very high colony counts (up to ∼90%) for some of cell populations sorted from the BONCAT permafrost incubations (**Figure 5**). Colony morphology/color/size confirmed that some of the flow cytometry profiles/sorting gates were enriched in specific types of bacteria, some at very low abundance in the global community, identified following rRNA gene amplification and Sanger sequencing. Overall, using BONCAT-Live after permafrost incubations, we isolated two dozen distinct strains of bacteria, representing five phyla and including potential novel species based on rRNA similarity to described species (**Figure 5**). Several known psychrophiles (*Cryobacterium*, *Psychromonas*) were recovered primarily after plating at 15°C and displayed impaired growth at 25°C. Most of the other isolates grew faster at 25°C. Using standard versus diluted nutrient agar did not appear to influence the types of recovered strains.

### BONCAT-Live in oral microbiota

As both rhizosphere soil and permafrost represent open environments, we also aimed to test the applicability of BONCAT-Live in a more selective host-microbiota system, the human microbiome. Mammalian-associated microbes have adapted to specialized niches, and many rely on cross-feeding or host-supplied nutrients and co-factors. We selected the oral environment, as it encompasses a multitude of nutritional niches that harbor high taxonomic diversity and a combination of facultative and strict anaerobes. BONCAT has already been demonstrated as a feasible approach in studying oral microbiota (15). Here, we investigated several nutrients that support and modulate interspecies interactions in saliva and supragingival biofilm. Because many oral bacteria are anaerobes, we performed all incubations, click-labeling and flow cytometry under anoxic conditions. In addition, due to the generally fast growth rate and to avoid extensive interspecies cross-feeding, we used a shorter incubation time than for environmental samples, 2 hours.

Collectively, the oral microbiota has complex nutritional requirements that may compete with the uptake/incorporation of HPG by some of its members. Therefore, to evaluate the breadth of BONCAT coverage under our experimental conditions, we first performed incubations in a non-selective, nutrient rich medium (MTGE) that supports the growth of many oral fastidious bacteria. Out of the 69 bacterial genera detected in the starting oral sample by amplicon sequencing, 52 were present in the BONCAT-labeled sorted population (**Figure 6** and **Supplemental Table**). Notable taxa that increased in relative abundance, based on both DNA staining and BONCAT, included *Gemella, Peptostreptococcus, Veilonella, Leptotrichia* and *Aggregatibacter*, while some displayed lower levels (*Actinomyces, Prevotella*). In particular, *Leptotrichia* (Fusobacterota) and *Streptococcus* (Firmicutes), had the strongest fluorescence signal, suggesting high translational activity.

**Figure 6.**
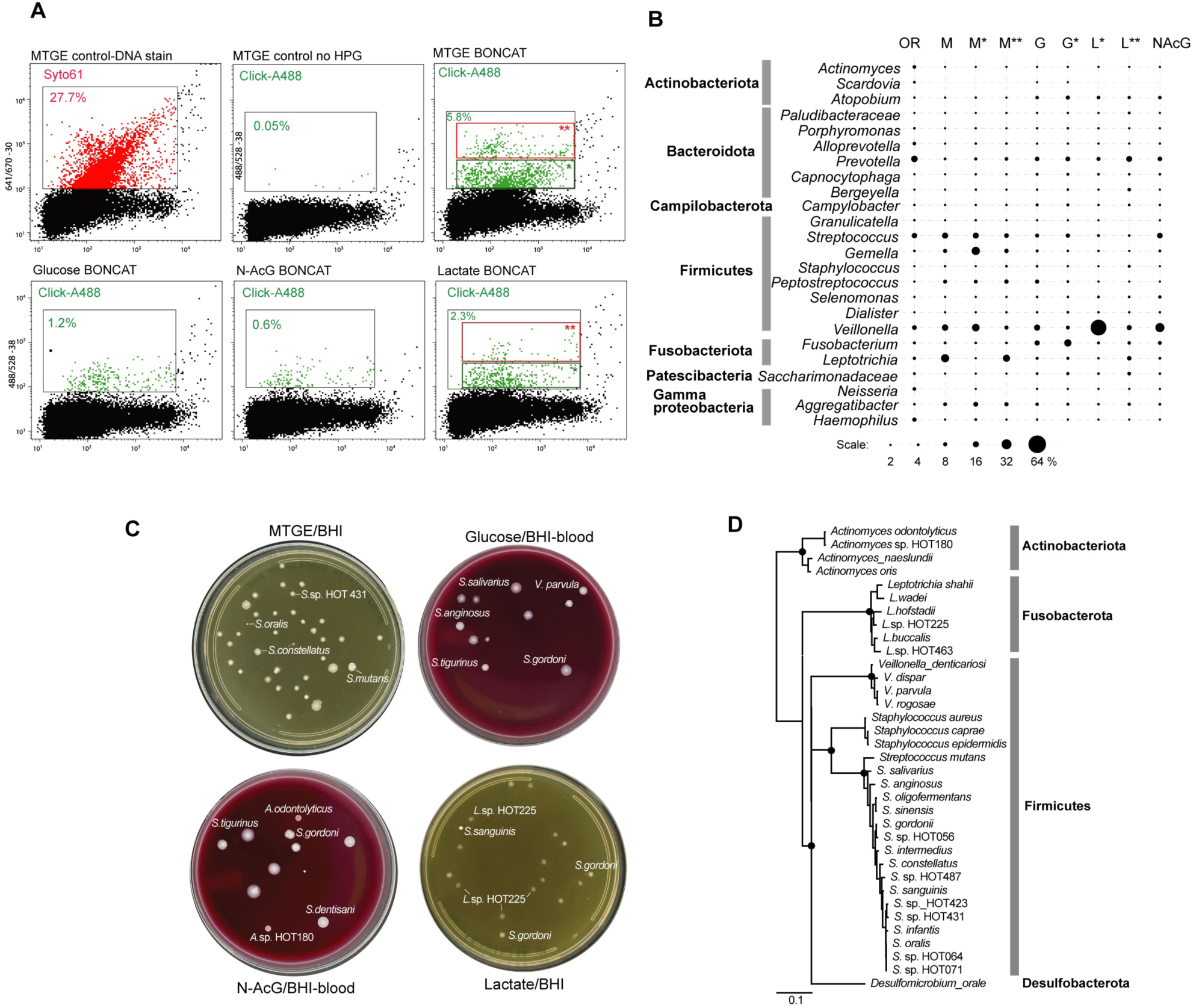
Oral microbiota BONCAT-Live. **(A)** Flow cytometry of human oral microbiota (saliva/supragingival biofilm sample) labeled after BONCAT incubation in MTGE medium or in basal medium supplemented with glucose, N-acetylglucosamine or lactate. The different gates used for sorting and particle/cell frequency are indicated. **(B)** Microbial taxa relative abundance (top genera) in original sample (OR) and BONCAT-positive sorted cell populations based on gating shown in (A). **(C)** Bacterial isolates following BONCAT-positive single cell sorting on BHI nutrient agar. Identity of representative colonies is indicated. **(D)** SSU rRNA phylogeny of isolated bacterial strains based on closest human oral relatives.

We then used a medium base with a lower nutrient content (artificial saliva), to determine how individual bacteria respond to the addition of selected growth substrates. Glucose and other sugars impact the metabolism of a variety of oral bacteria and, through acidogenic effects and community changes in the dental biofilm, are linked to carries (59). Based on BONCAT signal, the most active taxa upon being provided glucose were *Atopobium* (Actinobacteriota), *Prevotella* (Bacteroidota), *Streptococcus* and *Fusobacterium*. Lactate, a product of carbohydrate metabolism is also utilized by many oral species. The transition from sugars to lactate and organic acids utilization has been well documented, including by functional genomics (60). Lactate-containing media led to highest activity by *Prevotella, Veilonella* and *Leptotrichia*, while *Streptococcus* subsided (**Figure 6** and **Supplemental Table**). The third substrate we tested, N-acetyl glucosamine (N-AcGlc), is an important component of the extracellular matrix, mucins and bacterial cell walls. While some species recycle such amino sugars as peptidoglycan building blocks, nutritional utilization supports a diverse community through spatial, physiological niche partitioning and competition, e.g. between species of *Streptococcus* (61–64). In our incubations, N-AcGlc had the smallest effect in inducing high translational activity (BONCAT signal), presumably due to a slower rate of uptake and a more restricted utilization as compared to the other tested nutrients. Nevertheless, *Streptococcus* and *Veilonella* exhibited strong signal, corroborating their known metabolic and physical interaction within the community (60, 64, 65).

Single cells from the BONCAT positive populations identified by flow cytometry were deposited on non-selective nutrient agar and incubated anaerobically until colonies formed. The near full length 16S rRNA amplicons enabled assignments of those isolates to known human oral bacteria species/strains. Across all tested nutrients, described above, we isolated and assigned strains to over 30 human oral bacteria (**Figure 6B**). Expectedly, most colonies represented a variety of strains of *Streptococcus*, *Leptotrichia* and *Veilonella*, which were identified as highly abundant and active based on BONCAT population-level amplicon data. We also isolated representatives of *Actinomyces* and *Staphylococcus*, which were in the minority of BONCAT responders. On the other hand, we did not retrieve some other relatively abundant BONCAT-positive bacteria, including *Prevotella*, *Gemella* and *Fusobacterium*. This may have been due to inhibition caused by BONCAT/flow sorting or non-permissive culture conditions for those strains, which we did not attempt to distinguish and address here.

## DISCUSSION

Traditional and high throughput cultivation strategies have expanded the diversity of cultured organisms, but they do not directly or in real time inform the range of physiological aptitudes of those isolates and how they interact with a host, other microbes, or the environment, in a native setting. Cultivation-independent approaches have been designed to address such *in situ*, including stable isotope probing (SIP) and various meta-omics (10, 11, 66–68). BONCAT and other click-chemistry enabled cellular labeling techniques complement sequence-based interrogations of active microbes in their environmental settings. Combined with advanced spectrometry imaging techniques (nano-SIMS, Raman), isotope probing and BONCAT enable cell-level resolution assignment of metabolic and biogeochemical processes (69–71). Ideally, cells labeled based on their physiological properties should not only be identified destructively but retrieved viably for cultivation, for further characterization and downstream applications.

BONCAT-Live directly enables isolation and cultivation of microbial strains that are physiologically active or activated under specific conditions on environmentally relevant timeframes. Because the initial “growth” (protein synthesis that enables cell labeling) can occur in the native sample, the cells are exposed to some of their naturally occurring physical, chemical and biological factors. In two of the test cases we analyzed, rhizosphere and permafrost, samples were handled and incubated under conditions to maintain their native architecture as much as possible, so that microbial communities are primarily responding to the testing variables. Because the oral microbiota could not be studied *in vivo*, we tested responses to chemical cues under laboratory conditions permissive to some of the community members. Most of the relatively abundant BONCAT-positive taxa were retrieved as isolates, and the near full length 16S rRNAs resolved a higher strain diversity than attainable by short amplicons (e.g. the permafrost *Cryobacterium* and *Arthrobacter*, the rhizosphere *Pseudomonas*, the oral *Streptococcus* and *Leptotrichia,* **Figures 4, 5, 6**). Applying multiple sorting gates (**Figures 5, 6**) also led to isolating some low abundance positive taxa, half of the diversity of isolates being present at less than 1% abundance in the BONCAT-positive population. Therefore, while not a quantitative assay, single cell isolation and cultivation by BONCAT-Live can provide more confidence of microbial activity of individual taxa under specific conditions than sequences retrieved by cultivation-independent amplicon or metagenome sequencing. Follow up characterization of such strains would be necessary to test their physiological properties and how they relate to the BONCAT experiment variables or the biological/environmental questions being tested. For example, when investigating metabolites/compounds that can act as inhibitors, physiologically active microbes may be resistant to their effects but utilize other existing nutrients or use them as nutrients, either independently or in association with other members of the community through cross-feeding. Salicin is one such example we tested in the rhizosphere. The rapid, sharp increase in *Pseudomonas* and the *Micrococcaceae* actinobacteria (**Figure 3**), both metabolically versatile taxa, suggests they may have utilized salicin directly. A variety of other taxa proliferated as well as compared to the native rhizosphere incubation, leading to a nearly fourfold increase in the frequency of BONCAT-positive cells. Salicin utilization assays would be required to identify which of those isolates can uptake it and which use the salicyl alcohol resulting from salicin hydrolysis by other strains (43). The nature of the substrate and the selected timeframe are therefore important considerations in designing the BONCAT assay to probe the nutritional dynamics and to separate primary users from secondary cross-feeders.

Because taxa retrieval in BONCAT-Live is not quantitative, time course amplicon data of BONCAT-positive sorted populations could indicate such microbial succession and guide which isolates may correspond to which category, subject to physiological testing. The dynamics of the community and succession of various members can be approached by BONCAT and BONCAT-Live under a variety of settings and time frames, including *in situ* (19). Each of the technical steps of BONCAT-Live can affect the viability of individual taxa differently, which constitutes a limitation of the approach. Copper was, in our testing, the most problematic and, while reducing its concentration and the exposure to oxygen radicals improved viability of test bacteria and enabled isolation of a wide variety of other taxa, it is conceivable some microbes may not tolerate. While Cu-independent click chemistry labeling is a complementary option to be further explored, it introduces additional chemistries and incubation conditions that may also be detrimental to some microbes. Because BONCAT-Live provides the initial single cell isolation step from its native community, it is exposed to some of the same shortfalls of standard isolation by plating, namely media selection, incubation conditions, which would have to be explored and optimized depending on samples and target organisms. For example, selected single cells could be deposited on plates or in liquid culture that harbor helper organisms or grown as mini-communities by co-sorting from the BONCAT-positive population. Magnetic beads-based enrichments, demonstrated in our test experiments, could be an alternative to retrieve and culture population of microbes that are co-dependent, refractory to flow sorting or, by performing all steps in an anaerobic chamber, avoid toxicity for samples highly sensitive to oxygen exposure.

By broadly targeting physiologically (translationally) active microbes, BONCAT-Live is taxon-agnostic. Taxonomic identification of bacteria and archaea has been previously linked to BONCAT using FISH with specific probes following fixation (21, 69). Taxon-specific cell surface antibodies may be combined with and used to isolate specific taxa by BONCAT-Live by targeted reverse genomics (72). BONCAT-Live and other approaches to isolate and culture metabolically active single cells (73) therefore complement culture-independent strategies to characterize environmental microbes, communities and processes. BONCAT-Live does not only enable selecting a metabolic characteristic as the driver for isolation but combined with FACS fractionation provides access to low abundance taxa. The bacterial strains responding to specific stimuli that we isolated from the various environments will serve for further physiological characterization, individually and as part of synthetic communities.

## MATERIALS AND METHODS

### Environmental samples

#### Arctic samples

In March 2021, 8 boreholes were drilled at Ny-Ålesund, Svalbard, Norway, at the Bayelva long term permafrost monitoring site (78.92094° N, 11.83334° E) (54). For this study we used a core collected at borehole 7 (BH7), where the permafrost underlies the active layer at about 1.5 m and below. The soil is silty clay to sandy silt and is characterized as high lithic and low nutrient (58). The cores were shipped frozen and were stored at −20°C.

#### Rhizosphere samples

Near-surface (2-10 inches deep) clusters of fine roots (<3 mm diameter) and associated soil were collected with a trowel from 4-5 years old *Populus deltoides* trees at an experimental site on the ORNL reservation in Oak Ridge, TN, USA during active growth season (April-October). The soil and roots were transported in closed sterile bags to prevent drying in an insulated box to the laboratory and used within 3 days of collection.

#### Oral samples

Oral samples were provided by adult volunteers, self-declared to be orally healthy, in accordance with a protocol approved by the Oak Ridge Site-Wide Institutional Review Board (FWA 00005031)(74). Supragingival and gum line plaque was self-collected from premolars-molars on each mouth side using two OMR-110 swabs (DNA Genotek, Ottawa, Canada). The swab tips were immediately placed in a vial of liquid dental transport medium (LDT, Anaerobe Systems, Morgan Hill, CA) and processed within 2 hours.

Aliquots of all collected samples were stored frozen at −80°C for baseline microbial community characterization.

### BONCAT incubations

#### Cultured bacteria strains

For control tests and optimization experiments we used previously isolated *Populus* rhizosphere soil strains (*Pseudomonas* sp. GM41, *Terriglobus* sp. ORNL, *Flavobacterium* sp. CF108, *Bacillus* sp. OV166, *Rhizobium* sp. CF142, *Paraburkholderia* sp BT03. and *Roseimicrobium* sp. ORNL1)(30, 75, 76) and Svalbard permafrost *Pseudomonas* sp. (57). The strains were maintained individually in R2A medium (Thermo Scientific) at 25°C. For BONCAT incubations, we used a low methionine version of R2A (based on DSMZ medium 830 formulation) in which we substituted the peptone and casamino acids with 1 g/l of a synthetic, equal mix of all other 19 amino acids (referred to as defined R2A or R2A*). Each strain was inoculated in 5 ml R2A* with or without 50μM HPG (Vector Labs, Newman CA) and incubated for 48 hours at 25°C. For killed incorporation controls, cultures fixed with 3% paraformaldehyde were supplemented with HPG and incubated similarly. At the end of incubations, cells were collected by centrifugation (10 min at 8000g), washed three times in PBS and resuspended in PBS. Aliquots were stored in PBS containing 10% glycerol and 1% trehalose at −80°C or used immediately for click-chemistry labeling.

#### Rhizosphere incubations

Fragmented roots and associated rhizoplane soil (5 g) were spread on the surface of Petri plates and soaked with 10 ml 0.25 X Hoagland’s No 2 hydroponic mineral nutrient solution (Sigma-Aldrich, St. Louis MO) at 25°C for 18 hours. An additional 10 ml supplemented with 100 μM HPG was added, or supplement also containing 40 mM salicin (Sigma-Aldrich), or (40 mM) potassium malate (Sigma-Aldrich). Background control incubations contained no HPG or paraformaldehyde (3%). After gentle mixing, the plates were incubated for an additional 24 hours at 25°C. The roots, soil slurry and incubation solution were transferred into 50 ml Falcon tubes, supplemented with 10 ml PBS, vortexed for 1 minute and centrifuged at 1000g for 2 minutes. The supernatant was then filtered through a 50 microns CellTrics strainer (Sysmex, Lincolnshire, IL) and spun at 8000g for 10 minutes to pellet the microbial cells. The pellet was washed three times as above. Sample aliquots were used immediately for click-chemistry labeling and flow cytometry cell sorting and others preserved at −80°C in PBS-glycerol-trehalose.

#### Arctic soil incubations

Frozen soil samples (∼15 g) from active layer and permafrost horizons of the cores were distributed in 50 ml vented, 0.22 μm filter-capped Falcon tubes on ice. 7 ml ice-cold sterile 50 uM HPG solution in water was added to penetrate the soil. Control incubation samples received only water or HPG in 3% paraformaldehyde (killed control). To simulate permafrost locations with higher organic matter content, a set of incubations was supplemented with 10% R2A*. Incubations were performed at 4°C (2-6 weeks) or at −20°C (12 weeks). Following incubation, the samples were processed as described for the rhizosphere.

#### Oral microbiome incubations

To maintain viability of anaerobic bacteria, all oral sample processing was performed under anoxic conditions in a COY anaerobic chamber with an atmosphere of 85% N_2_, 10% CO_2_ and 5% H_2_. The vial with oral swabs in LDT were transferred into the chamber into and vortexed for 1 minute. The liquid was passed through a 10 μm CellTrics strainer into Eppendorf tubes, centrifuged for 10 minutes at 8000g and the pellet washed and resuspended in 2 ml PBS. Aliquots (100 μl) were added into 1 ml BONCAT incubation media with 100 μM HPG that included rich, non-selective MTGE medium (Anaerobe Systems) or chemically defined artificial saliva medium (DMM)(77) supplemented with either 10 mM glucose, 10 mM sodium lactate or 10 mM N-acetyl glucosamine. As controls, samples were inoculated in MTGE medium with no HPG or with 3% paraformaldehyde. After incubation for 2 hours at 37°C, the samples were centrifuged, washed three times with PBS and immediately used for click chemistry labeling.

### Click chemistry biotinylation for imaging and magnetic enrichment

Six of the cultured test bacteria (*Terriglobus*, *Flavobacterium*, *Bacillus*, *Rhizobium*, *Paraburkholderia* and *Roseimicrobium*) grown in the presence of HPG and their corresponding negative controls (killed and non-HPG) were used individually for in suspension labeling experiments guided by a published protocol (78). Freshly prepared aminoguanidine hydrochloride (50 μl, 100 mM in PBS) and sodium ascorbate (50 μl, 100 mM in PBS) were added to 0.9 ml cell suspension (OD_600_=1.0) in PBS in a 1.5 ml Eppendorf tube and mixed by inversion. A separately prepared mix containing CuSO_4_ (5 μl, 20 mM in water), Tris[(1-hydroxypropyl-1H-1,2,3-triazol-4-yl)methyl]amine (THPTA, Click Chemistry Tools) (10 μl, 50 mM in water) and biotin picolyl azide (Click Chemistry Tools) (5 μl, 2 mM in DMSO), pre-reacted for 3 minutes, was pipetted in and the tube inverted once. Following 30 minutes incubation in the dark, cells were pelleted washed three times in PBS and resuspended in 1 ml PBS. For imaging, 50 μl aliquots were mixed with a 50 μl PBS solution containing 1% biotin-free BSA (Fraction 5, Sigma), 0.02% Pluronic F127 surfactant (Sigma) and 5 μg/ml streptavidin-Alexa Fluor 488 conjugate (ThermoFisher), incubated in the dark for 30 minutes, washed three times with PBS-0.02% Pluronic F127 and visualized and photographed under epifluorescence on a Zeiss AxioImager M2 microscope. Because streptavidin cannot penetrate unfixed cells, fluorescent labeling was at the cell surface level. No staining was observed for negative control incubations.

For the magnetic enrichment experiment, we obtained a purified microbial fraction from 10 grams of rhizosphere soil from the *Populus* plantation using centrifugation on Histodenz, as we described previously (79). Approximately 5×10^8^ rhizosphere bacteria (based on an estimation of 1OD_600_=10^9^ cells) was mixed with 10^8^ of each individual biotinylated strain and adjusted to 1 ml in PBS. An aliquot (50 μl) was saved for community diversity characterization and the rest was mixed with 1 ml PBS containing 1 mg/ml biotin-free BSA, 0.02% Pluronic F127 and 2 mM EDTA (PBS*). We then added 200 μl Streptavidin MicroBeads (Miltenyi Biotech, Gaithersburg, MD) and incubated the suspension at 4°C for two hours on a rotator. The cells-beads suspension was then pelleted by centrifugation (8000g, 10 minutes at 4°C), washed twice with 1 ml PBS* and resuspended in 0.5 ml PBS*. An LS MACS column and separator (Miltenyi Biotech) was then used, following the manufacturers protocol, to purify the magnetically labeled biotinylated cells. The purified cell fraction, along with aliquots of original soil community and the soil-test strains mix were used for DNA extraction with the ZymoBIOMICS DNA Microprep Kit (Zymo Research, Irvine CA).

### Viability tests and click chemistry labeling for BONCAT-Live

To test the impact of various chemical compounds on viability we used the soil rhizosphere strain *Pseudomonas* sp. GM41, the output being colony formation after single cell flow sorting on nutrient agar. Each of the major chemicals used in click-chemistry reactions (sodium ascorbate, aminoguanidine hydrochloride, 2-chloroacetamide, CuSO4, THPTA, AZDye 488 DBCO or AZDye488 Picolyl Azide (Click Chemistry Tools) was individually tested at the concentration recommended for click labeling reactions (78) by incubation with cells suspended in PBS at room temperature for 30 minutes (37°C for chloroacetamide), followed by cell washing in PBS and flow sorting. Parallel tests were conducted with the complete click-labeling reaction protocol, as described above and in (78). The only major negative impact was observed for 2-chloroacetamide (∼80% inactivation) and CuSO_4_ at 50-100 μM (>50% inactivation). Consequently, we did not further apply the strain-promoted (Cu-free) click chemistry protocol and we focused on improving the viability with Cu-catalyzed reactions. Three combined modifications to the standard protocol restored viability (>80%) while maintaining effective fluorescent labelling with 2 μM AZDye488 Picolyl Azide. First, CuSO_4_ concentration was reduced to 5 μM. Second, we substituted THPTA with BTTAA (Click Chemistry Tools), an improved reaction accelerator that also has lower cytotoxicity (80). Third, we added to the cell suspension (prior to introduction of reaction chemicals), 1 mM sodium pyruvate (Sigma) and 100 U/μl catalase (Bovine liver, Sigma), reactive oxygen scavengers that improve microbial viability (36, 38, 81). These were then applied to fluorescent labeling of microbiota samples following BONCAT incubations. For click reactions with oxygen sensitive oral microbiota samples, we prepared all fresh stock solutions and degassed buffer under nitrogen gas bubbling with long needles inserted in the Falcon tubes followed by capping and transfer to anaerobic chamber.

For microbiota samples that we used in BONCAT-Live sorting experiments, samples were mostly labeled fresh, immediately after BONCAT incubations. We also labeled and sorted for cultivation soil samples that were cryopreserved (primarily from permafrost incubations). While it is likely not all microbes will survive freezing, we did not collect sufficient isolates to distinguish differences. Because immediate isolation is not always feasible after BONCAT, click labeling and FACS is applicable to preserved samples as well.

### Flow cytometry sorting and cultivation

Prior to flow cytometry analyses, pure cultures used for viability testing and microbiota samples were counterstained for DNA with 0.5 μM Syto61 red fluorescent dye (ThermoFisher Scientific). For flow cytometry and cell sorting, we used a Cytopeia/BD Influx Model 208S (BD, San Jose, CA). The sheath fluid was 0.5xPBS (filtered through 0.1 μm filter), pressurized at 18 psi. For cytometry analyses and sorting, we used a 70 μm nozzle and the 488 nm laser for forward-side scatter (FSC-SSC) analyses and the 488 nm and 641 nm lasers for fluorescence detection and cell sorting trigger. Gating parameters were selected based on FSC-SSC and red/green fluoresce levels for each experiment and sample to include DNA staining and, when click labeled with Alexa488, BONCAT surface staining. For sorting pools of cells from selected gated populations 5,000-20,000 cells were deposited per wells containing 3 ml sterile, UV-irradiated TE (82) in a 96 well plate (10mM Tris-HCl pH 8.0, 1 mM EDTA). In our workflow, a minimum of 5,000 sorted bacterial cells were determined necessary for reliable generation of rRNA amplicon libraries for sequencing. The pools of cells were stored at −80°C after sealing with sterile foil.

To sort single cells for cultivation, we used standard round or rectangular culture plates with selected nutrient media agar for the different microbiota samples. The sorting chamber was sterilized by UV with the instrument germicidal lamp for 15 minutes. Deposition of 100 cells per plate was performed with sorting set to single particle pure mode. After sorting, plates were incubated at 4°C (permafrost, 7+ days), 25°C (rhizosphere and permafrost, 24-72 hours). For experiments with oral microbiota samples we prepared the instrument for anoxic sorting, as previously described (25). Briefly, the sheath fluid was heated to 95°C in the pressure vessel and cooled under nitrogen. The flow sorter gas lines were switched to nitrogen. The oral samples in sorting tubes, prepared and capped in the anaerobic chamber, were inserted in the instrument sample port and pressurized with nitrogen gas. Following single cell deposition of oral bacteria on anoxic culture media, the plates were immediately transferred into acrylic boxes with oxygen-absorbing AnaeroPacks (Mitsubishi Gas Company) and incubated at 37°C (24-72 hours).

### Molecular microbial characterization

DNA was extracted from aliquots of starting environmental samples with the ZymoBIOMICS DNA Microprep Kit. Sorted pools of cells were subject to lysis by three cycles of snap freeze-heat between a metal 96 well block on dry ice followed by a PCR block at 95°C (3 minutes each). DNA and cell lysates were then used for microbial community characterization by small subunit (SSU) rRNA gene amplification (V4 region) using Illumina-adapted, barcoded universal primers 515F and 806R (detailed in (83)), with the Quick-16S NGS Library Prep Kit (Zymo Research), according to the manufacturer’s protocol. Bidirectional amplicon sequencing was performed on a MiSeq instrument (Illumina, San Diego, CA, USA) using the v2 500 cycles kit, according to manufacturer’s instructions. Demultiplexed raw sequence reads were imported into Qiime2 (84) and analyzed for taxonomic composition and relative abundance using standard workflow as described in (74, 85, 86). Bubble plots representing the most relatively abundant taxa were generated using a perl script (87).

To identify the microbial isolates, near full length SSU rRNA gene amplification was performed by colony PCR with bacterial universal primers (27F-1492R), followed by Sanger sequencing (Eurofin Genomics, Louisville KY). The sequences were imported in Geneious Prime v.2021 (88), trimmed based on quality and compared to relatives (cultured and uncultured) in Genbank using blastn. For the oral isolates, comparisons were done using the Human Oral Microbiome Database (https://www.homd.org/)(89). Sequences of our isolates and their closest relatives were aligned in Geneious using MUSCLE (90) and used for phylogenetic tree reconstruction with FastTree (91). The figures were composed and labeled in Adobe Illustrator.

## Materials and data availability

Bacterial isolates are available upon requests. The Sanger sequence data for the isolates has been deposited in GenBank under accession numbers XXXX.

## Acknowledgments

This research was sponsored by the Genomic Science Program, U.S. Department of Energy, Office of Science, Biological and Environmental Research, as part of the Plant Microbe Interfaces Scientific Focus Area (http://pmi.ornl.gov). Oak Ridge National Laboratory is managed by UT-Battelle, LLC, for the U.S. Department of Energy under contract DE-AC05-00OR22725. M.P. was also sponsored by grant R01DE024463 from the National Institute of Dental and Craniofacial Research of the U.S. National Institutes of Health. K.G.L. and T.A.V. were sponsored by grant DE-SC0020369 from the Genomic Science Program of the U.S. Department of Energy, Office of Science, Office of Biological and Environmental Research. S.A.M. was also sponsored by a UT Science Alliance Graduate Advancement Training and Education (GATE) program fellowship and by a University of Tennessee Faculty-Student Research Award with K.G.L.

## Notes

### Competing Interest Statement

The authors have declared no competing interest.

