## Supplemental Figure S1 for "BONCAT-Live for isolation and cultivation of active environmental microbes"

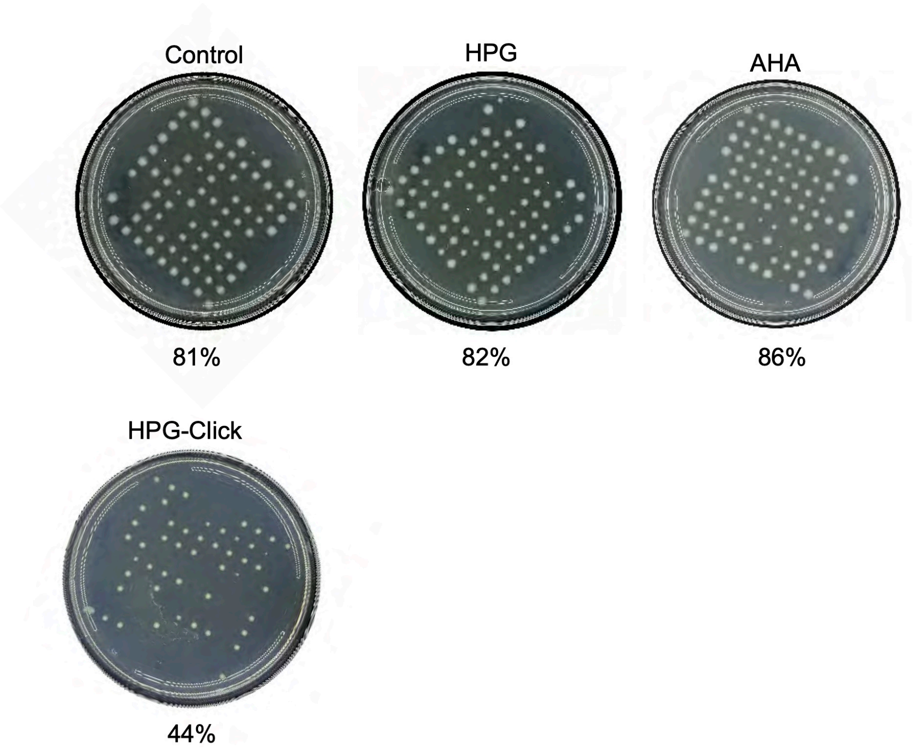


**Figure S1**. Viability tests for flow sorted *Pseudomonas* sp GH41. On each test, 100 particles were individually deposited on R2A agar plate, followed by incubation (2 days at 25C). Control, untreated cells grown in R2A*; HPG and AHA, cells grown in R2A* supplemented with HPG or AHA, respectively; HPG-Click, cells grown in R2A* supplemented with HPG that were click-labeled using copper chemistry protocol. The percentages indicate number of colonies.
