## Supplemental tables for "BONCAT-Live for isolation and cultivation of active environmental microbes"

**Relative taxon abundance in rhizosphere soil and in populations sorted follwing BONCAT incubations.**

R, sorted based on red fluoresence (DNA stain). G, sorted based on A488 click label

| Taxon | Rhizosphere | NoHPG R | HPG R | HPG Low G | HPG High G | HPG Salicin R | HPG Salicin G | HPG Malate R | HPG Malate G |
|---|---|---|---|---|---|---|---|---|---|
| Acidobacteriota;c__Acidobacteriae;o__Acidobacteriales | 0.18 | 0.00 | 0.00 | 0.00 | 0.00 | 0.00 | 0.00 | 0.00 | 0.00 |
| Acidobacteriota;c__Acidobacteriae;o__Acidobacteriales;f__Acidobacteriaceae_(Subgroup_1) | 0.13 | 0.18 | 0.10 | 0.01 | 0.00 | 0.00 | 0.01 | 0.00 | 0.00 |
| Acidobacteriota;c__Acidobacteriae;o__Bryobacterales;f__Bryobacteraceae;g__Bryobacter | 0.32 | 0.00 | 0.00 | 0.00 | 0.00 | 0.00 | 0.00 | 0.00 | 0.00 |
| Acidobacteriota;c__Acidobacteriae;o__Solibacterales;f__Solibacteraceae;g__Candidatus_Solibacter | 0.84 | 0.00 | 0.00 | 0.02 | 0.00 | 0.00 | 0.00 | 0.00 | 0.00 |
| Acidobacteriota;c__Acidobacteriae;o__Subgroup_2;f__Subgroup_2;g__Subgroup_2 | 0.37 | 0.00 | 0.00 | 0.00 | 0.00 | 0.00 | 0.00 | 0.00 | 0.00 |
| Acidobacteriota;c__Blastocatellia;o__11-24;f__11-24;g__11-24 | 0.69 | 0.00 | 0.00 | 0.00 | 0.00 | 0.00 | 0.00 | 0.00 | 0.00 |
| Acidobacteriota;c__Blastocatellia;o__Blastocatellales;f__Blastocatellaceae | 0.17 | 0.00 | 0.00 | 0.00 | 0.00 | 0.00 | 0.00 | 0.00 | 0.00 |
| Acidobacteriota;c__Blastocatellia;o__Elev-16S-573;f__Elev-16S-573;g__Elev-16S-573 | 0.10 | 0.00 | 0.00 | 0.00 | 0.00 | 0.00 | 0.00 | 0.00 | 0.00 |
| Acidobacteriota;c__Blastocatellia;o__Pyrinomonadales;f__Pyrinomonadaceae;g__RB41 | 1.19 | 0.00 | 0.00 | 0.00 | 0.00 | 0.00 | 0.00 | 0.00 | 0.00 |
| Acidobacteriota;c__Holophagae;o__Subgroup_7;f__Subgroup_7;g__Subgroup_7 | 0.54 | 0.00 | 0.00 | 0.00 | 0.00 | 0.00 | 0.00 | 0.00 | 0.00 |
| Acidobacteriota;c__Subgroup_22;o__Subgroup_22;f__Subgroup_22;g__Subgroup_22 | 0.19 | 0.00 | 0.00 | 0.00 | 0.00 | 0.00 | 0.00 | 0.00 | 0.00 |
| Acidobacteriota;c__Subgroup_25;o__Subgroup_25;f__Subgroup_25;g__Subgroup_25 | 0.13 | 0.00 | 0.00 | 0.00 | 0.00 | 0.00 | 0.00 | 0.00 | 0.00 |
| Acidobacteriota;c__Subgroup_5;o__Subgroup_5;f__Subgroup_5;g__Subgroup_5 | 0.28 | 0.00 | 0.00 | 0.00 | 0.00 | 0.00 | 0.00 | 0.00 | 0.00 |
| Acidobacteriota;c__Vicinamibacteria;o__Subgroup_17;f__Subgroup_17;g__Subgroup_17 | 0.79 | 0.00 | 0.05 | 0.00 | 0.00 | 0.04 | 0.00 | 0.03 | 0.00 |
| Acidobacteriota;c__Vicinamibacteria;o__Vicinamibacterales;f__uncultured;g__uncultured | 7.52 | 0.31 | 0.04 | 0.03 | 0.07 | 0.20 | 0.00 | 0.01 | 0.02 |
| Acidobacteriota;c__Vicinamibacteria;o__Vicinamibacterales;f__Vicinamibacteraceae;g__uncultured | 0.21 | 0.00 | 0.00 | 0.00 | 0.00 | 0.00 | 0.00 | 0.00 | 0.00 |
| Acidobacteriota;c__Vicinamibacteria;o__Vicinamibacterales;f__Vicinamibacteraceae;g__Vicinamibacteraceae | 1.13 | 0.13 | 0.06 | 0.00 | 0.00 | 0.05 | 0.00 | 0.00 | 0.01 |
| Actinobacteriota;c__Acidimicrobiia;o__IMCC26256;f__IMCC26256;g__IMCC26256 | 1.60 | 0.00 | 0.00 | 0.00 | 0.00 | 0.00 | 0.00 | 0.00 | 0.00 |
| Actinobacteriota;c__Acidimicrobiia;o__Microtrichales | 0.19 | 0.00 | 0.00 | 0.00 | 0.00 | 0.00 | 0.00 | 0.00 | 0.02 |
| Actinobacteriota;c__Acidimicrobiia;o__Microtrichales;f__Ilumatobacteraceae | 0.12 | 0.05 | 0.00 | 0.00 | 0.00 | 0.00 | 0.00 | 0.00 | 0.00 |
| Actinobacteriota;c__Acidimicrobiia;o__Microtrichales;f__Ilumatobacteraceae;g__CL500-29_marine_group | 0.23 | 0.00 | 0.02 | 0.00 | 0.00 | 0.04 | 0.00 | 0.01 | 0.00 |
| Actinobacteriota;c__Acidimicrobiia;o__Microtrichales;f__Ilumatobacteraceae;g__uncultured | 0.58 | 0.00 | 0.00 | 0.00 | 0.00 | 0.00 | 0.00 | 0.00 | 0.00 |
| Actinobacteriota;c__Acidimicrobiia;o__Microtrichales;f__uncultured;g__uncultured | 0.32 | 0.00 | 0.00 | 0.00 | 0.00 | 0.00 | 0.00 | 0.00 | 0.00 |
| Actinobacteriota;c__Actinobacteria | 1.41 | 0.12 | 0.79 | 1.01 | 0.00 | 2.82 | 4.78 | 0.10 | 0.00 |
| Actinobacteriota;c__Actinobacteria;o__Corynebacteriales;f__Mycobacteriaceae;g__Mycobacterium | 0.65 | 0.27 | 0.38 | 2.12 | 0.00 | 0.09 | 0.32 | 0.06 | 0.04 |
| Actinobacteriota;c__Actinobacteria;o__Frankiales;f__Acidothermaceae;g__Acidothermus | 0.65 | 0.00 | 0.00 | 0.00 | 0.00 | 0.00 | 0.00 | 0.00 | 0.00 |
| Actinobacteriota;c__Actinobacteria;o__Frankiales;f__Frankiaceae;g__Jatrophihabitans | 0.22 | 0.00 | 0.00 | 0.00 | 0.00 | 0.00 | 0.00 | 0.00 | 0.00 |
| Actinobacteriota;c__Actinobacteria;o__Frankiales;f__Nakamurellaceae;g__Nakamurella | 0.04 | 0.00 | 0.00 | 0.00 | 0.00 | 0.09 | 0.13 | 0.00 | 0.00 |
| Actinobacteriota;c__Actinobacteria;o__Frankiales;f__Sporichthyaceae | 0.13 | 0.00 | 0.00 | 0.00 | 0.00 | 0.00 | 0.00 | 0.00 | 0.00 |
| Actinobacteriota;c__Actinobacteria;o__Micrococcales;f__Microbacteriaceae | 0.05 | 0.36 | 1.87 | 0.79 | 0.00 | 1.54 | 0.07 | 0.13 | 0.37 |
| Actinobacteriota;c__Actinobacteria;o__Micrococcales;f__Microbacteriaceae | 0.11 | 1.43 | 4.39 | 0.94 | 0.05 | 2.57 | 0.15 | 0.42 | 0.91 |
| Actinobacteriota;c__Actinobacteria;o__Micrococcales;f__Microbacteriaceae;g__Agromyces | 0.04 | 0.10 | 0.00 | 0.00 | 0.00 | 0.00 | 0.00 | 0.00 | 0.00 |
| Actinobacteriota;c__Actinobacteria;o__Micrococcales;f__Microbacteriaceae;g__Microbacterium | 0.02 | 0.97 | 2.52 | 0.15 | 0.05 | 1.03 | 0.07 | 0.29 | 0.54 |
| Actinobacteriota;c__Actinobacteria;o__Micrococcales;f__Micrococcaceae | 0.06 | 1.87 | 2.11 | 2.12 | 0.07 | 20.02 | 25.22 | 0.89 | 0.21 |
| Actinobacteriota;c__Actinobacteria;o__Micromonosporales;f__Micromonosporaceae | 0.65 | 0.02 | 0.03 | 0.00 | 0.03 | 0.00 | 0.00 | 0.00 | 0.00 |
| Actinobacteriota;c__Actinobacteria;o__Micromonosporales;f__Micromonosporaceae;g__Dactylosporangium | 0.11 | 0.00 | 0.00 | 0.00 | 0.00 | 0.00 | 0.00 | 0.00 | 0.00 |
| Actinobacteriota;c__Actinobacteria;o__Propionibacteriales;f__Nocardioidaceae | 0.29 | 0.20 | 0.00 | 0.00 | 0.00 | 0.39 | 0.05 | 0.00 | 0.00 |
| Actinobacteriota;c__Actinobacteria;o__Propionibacteriales;f__Nocardioidaceae | 1.42 | 0.75 | 2.93 | 0.16 | 0.08 | 1.31 | 0.31 | 0.34 | 0.08 |
| Actinobacteriota;c__Actinobacteria;o__Propionibacteriales;f__Nocardioidaceae;g__Aeromicrobium | 0.00 | 0.07 | 1.17 | 0.04 | 0.00 | 0.00 | 0.00 | 0.16 | 0.03 |
| Actinobacteriota;c__Actinobacteria;o__Propionibacteriales;f__Nocardioidaceae;g__Kribbella | 0.12 | 0.00 | 0.00 | 0.00 | 0.00 | 0.00 | 0.00 | 0.00 | 0.00 |
| Actinobacteriota;c__Actinobacteria;o__Propionibacteriales;f__Nocardioidaceae;g__Marmoricola | 0.04 | 0.05 | 0.04 | 0.00 | 0.00 | 0.00 | 0.00 | 0.00 | 0.00 |
| Actinobacteriota;c__Actinobacteria;o__Propionibacteriales;f__Nocardioidaceae;g__Nocardioides | 0.97 | 0.43 | 1.71 | 0.12 | 0.08 | 0.92 | 0.26 | 0.18 | 0.05 |
| Actinobacteriota;c__Actinobacteria;o__Propionibacteriales;f__Propionibacteriaceae;g__Microlunatus | 0.47 | 0.00 | 0.00 | 0.00 | 0.00 | 0.00 | 0.04 | 0.00 | 0.00 |
| Actinobacteriota;c__Actinobacteria;o__Pseudonocardiales;f__Pseudonocardiaceae;g__Pseudonocardia | 0.41 | 0.00 | 0.00 | 0.00 | 0.00 | 0.00 | 0.00 | 0.00 | 0.00 |
| Actinobacteriota;c__Actinobacteria;o__Streptomycetales;f__Streptomycetaceae | 0.24 | 0.00 | 0.00 | 0.00 | 0.07 | 0.35 | 0.65 | 0.00 | 0.00 |
| Actinobacteriota;c__Actinobacteria;o__Streptomycetales;f__Streptomycetaceae;g__Streptomyces | 0.27 | 0.00 | 0.00 | 0.00 | 0.00 | 0.00 | 0.00 | 0.00 | 0.00 |
| Actinobacteriota;c__Actinobacteria;o__Streptosporangiales;f__Thermomonosporaceae;g__Actinoallomurus | 0.13 | 0.00 | 0.00 | 0.00 | 0.00 | 0.00 | 0.00 | 0.00 | 0.00 |
| Actinobacteriota;c__MB-A2-108;o__MB-A2-108;f__MB-A2-108;g__MB-A2-108 | 1.97 | 0.00 | 0.00 | 0.00 | 0.00 | 0.00 | 0.00 | 0.00 | 0.00 |
| Actinobacteriota;c__Rubrobacteria;o__Rubrobacterales;f__Rubrobacteriaceae;g__Rubrobacter | 0.23 | 0.00 | 0.00 | 0.00 | 0.00 | 0.00 | 0.00 | 0.00 | 0.01 |
| Actinobacteriota;c__Thermoleophilia;o__Gaiellales | 4.51 | 0.02 | 0.01 | 0.00 | 0.00 | 0.00 | 0.00 | 0.00 | 0.00 |
| Actinobacteriota;c__Thermoleophilia;o__Gaiellales;f__Gaiellaceae;g__Gaiella | 1.31 | 0.00 | 0.00 | 0.00 | 0.00 | 0.11 | 0.00 | 0.00 | 0.00 |
| Actinobacteriota;c__Thermoleophilia;o__Gaiellales;f__uncultured;g__uncultured | 2.08 | 0.00 | 0.00 | 0.00 | 0.00 | 0.00 | 0.00 | 0.01 | 0.00 |
| Actinobacteriota;c__Thermoleophilia;o__Solirubrobacterales;f__67-14;g__67-14 | 2.48 | 0.03 | 0.00 | 0.00 | 0.00 | 0.00 | 0.00 | 0.00 | 0.02 |
| Actinobacteriota;c__Thermoleophilia;o__Solirubrobacterales;f__Solirubrobacteraceae | 0.63 | 0.00 | 0.00 | 0.00 | 0.00 | 0.00 | 0.00 | 0.00 | 0.00 |
| Actinobacteriota;c__Thermoleophilia;o__Solirubrobacterales;f__Solirubrobacteraceae;g__Conexibacter | 0.83 | 0.00 | 0.00 | 0.00 | 0.00 | 0.00 | 0.01 | 0.00 | 0.00 |
| Actinobacteriota;c__Thermoleophilia;o__Solirubrobacterales;f__Solirubrobacteraceae;g__Solirubrobacter | 0.29 | 0.00 | 0.00 | 0.00 | 0.00 | 0.13 | 0.00 | 0.00 | 0.00 |
| Actinobacteriota;c__Thermoleophilia;o__uncultured;f__uncultured;g__uncultured | 0.28 | 0.00 | 0.00 | 0.00 | 0.00 | 0.00 | 0.00 | 0.00 | 0.00 |
| Armatimonadota;c__Armatimonadia;o__Armatimonadales;f__Armatimonadales;g__Armatimonadales | 0.03 | 0.00 | 0.02 | 0.30 | 0.00 | 0.03 | 0.01 | 0.00 | 0.07 |
| Armatimonadota;c__Chthonomonadetes;o__Chthonomonadales;f__Chthonomonadaceae;g__Chthonomonas | 0.04 | 0.00 | 0.02 | 0.04 | 0.00 | 0.00 | 0.00 | 0.00 | 0.00 |
| Armatimonadota;c__Fimbriimonadia;o__Fimbriimonadales;f__Fimbriimonadaceae;g__Fimbriimonadaceae | 0.02 | 0.00 | 0.08 | 0.01 | 0.00 | 0.00 | 0.00 | 0.00 | 0.00 |
| Bacteroidota;c__Bacteroidia;o__Bacteroidales;f__Marinilabiliaceae;g__[Cytophaga]_xylanolytica_group | 0.00 | 0.00 | 0.00 | 0.00 | 0.00 | 0.00 | 0.00 | 0.00 | 0.00 |
| Bacteroidota;c__Bacteroidia;o__Chitinophagales;f__Chitinophagaceae | 0.35 | 0.66 | 0.17 | 0.01 | 0.04 | 0.09 | 0.72 | 0.20 | 0.02 |
| Bacteroidota;c__Bacteroidia;o__Chitinophagales;f__Chitinophagaceae | 0.63 | 0.72 | 0.72 | 0.48 | 0.06 | 0.40 | 1.30 | 0.64 | 0.97 |
| Bacteroidota;c__Bacteroidia;o__Chitinophagales;f__Chitinophagaceae;g__Aurantisolimonas | 0.02 | 0.00 | 0.00 | 0.00 | 0.00 | 0.00 | 0.03 | 0.00 | 0.21 |
| Bacteroidota;c__Bacteroidia;o__Chitinophagales;f__Chitinophagaceae;g__Chitinophaga | 0.01 | 0.01 | 0.00 | 0.22 | 0.00 | 0.21 | 0.08 | 0.00 | 0.46 |
| Bacteroidota;c__Bacteroidia;o__Chitinophagales;f__Chitinophagaceae;g__Edaphobaculum | 0.06 | 0.00 | 0.05 | 0.04 | 0.03 | 0.00 | 0.00 | 0.00 | 0.00 |
| Bacteroidota;c__Bacteroidia;o__Chitinophagales;f__Chitinophagaceae;g__Ferruginibacter | 0.07 | 0.00 | 0.00 | 0.01 | 0.00 | 0.00 | 0.07 | 0.00 | 0.02 |
| Bacteroidota;c__Bacteroidia;o__Chitinophagales;f__Chitinophagaceae;g__Heliimonas | 0.00 | 0.05 | 0.16 | 0.20 | 0.00 | 0.05 | 0.41 | 0.00 | 0.02 |
| Bacteroidota;c__Bacteroidia;o__Chitinophagales;f__Chitinophagaceae;g__Taibaiella | 0.00 | 0.00 | 0.34 | 0.00 | 0.00 | 0.04 | 0.00 | 0.44 | 0.24 |
| Bacteroidota;c__Bacteroidia;o__Chitinophagales;f__Chitinophagaceae;g__uncultured | 0.12 | 0.00 | 0.00 | 0.00 | 0.00 | 0.00 | 0.00 | 0.00 | 0.00 |
| Bacteroidota;c__Bacteroidia;o__Cytophagales;f__Spirosomaceae;g__Dyadobacter | 0.00 | 0.03 | 0.08 | 0.00 | 0.00 | 0.19 | 0.38 | 0.15 | 2.15 |
| Bacteroidota;c__Bacteroidia;o__Flavobacteriales;f__Flavobacteriaceae;g__Flavobacterium | 0.27 | 1.68 | 6.38 | 0.71 | 0.72 | 0.07 | 0.00 | 4.13 | 2.52 |
| Bacteroidota;c__Bacteroidia;o__Flavobacteriales;f__Weeksellaceae | 0.00 | 0.00 | 0.00 | 0.00 | 0.00 | 0.00 | 0.00 | 0.05 | 0.53 |
| Bacteroidota;c__Bacteroidia;o__Flavobacteriales;f__Weeksellaceae;g__Chryseobacterium | 0.00 | 0.49 | 0.64 | 0.23 | 0.09 | 0.35 | 0.13 | 1.38 | 0.76 |
| Bacteroidota;c__Bacteroidia;o__Sphingobacteriales;f__AKYH767;g__AKYH767 | 0.13 | 0.00 | 0.00 | 0.00 | 0.00 | 0.00 | 0.00 | 0.00 | 0.00 |
| Bacteroidota;c__Bacteroidia;o__Sphingobacteriales;f__Sphingobacteriaceae | 0.00 | 0.00 | 0.02 | 0.02 | 0.00 | 0.00 | 0.11 | 0.11 | 0.27 |
| Bacteroidota;c__Bacteroidia;o__Sphingobacteriales;f__Sphingobacteriaceae;g__Mucilaginibacter | 0.00 | 2.99 | 6.51 | 4.06 | 0.38 | 1.23 | 1.16 | 0.40 | 1.28 |
| Bacteroidota;c__Bacteroidia;o__Sphingobacteriales;f__Sphingobacteriaceae;g__Pedobacter | 0.00 | 0.80 | 3.93 | 0.67 | 0.12 | 0.28 | 0.15 | 4.10 | 3.27 |
| Bdellovibrionota;c__Bdellovibrionia;o__Bacteriovoracales;f__Bacteriovoracaceae;g__Bacteriovorax | 0.00 | 0.11 | 0.03 | 0.11 | 0.00 | 0.00 | 0.00 | 0.06 | 0.01 |
| Bdellovibrionota;c__Bdellovibrionia;o__Bdellovibrionales;f__Bdellovibrionaceae;g__Bdellovibrio | 0.10 | 0.00 | 0.00 | 0.00 | 0.00 | 0.00 | 0.00 | 0.00 | 0.00 |
| Bdellovibrionota;c__Oligoflexia;o__0319-6G20;f__0319-6G20;g__0319-6G20 | 0.18 | 0.00 | 0.00 | 0.00 | 0.00 | 0.00 | 0.00 | 0.00 | 0.00 |
| Chloroflexi;c__Anaerolineae;o__Anaerolineales;f__Anaerolineaceae;g__Anaerolinea | 0.26 | 0.00 | 0.00 | 0.00 | 0.00 | 0.00 | 0.00 | 0.00 | 0.00 |
| Chloroflexi;c__Anaerolineae;o__Anaerolineales;f__Anaerolineaceae;g__uncultured | 0.13 | 0.00 | 0.00 | 0.00 | 0.00 | 0.00 | 0.00 | 0.00 | 0.00 |
| Chloroflexi;c__Anaerolineae;o__Anaerolineales;f__Anaerolineaceae;g__UTCFX1 | 0.12 | 0.00 | 0.00 | 0.00 | 0.00 | 0.00 | 0.00 | 0.00 | 0.00 |
| Chloroflexi;c__Anaerolineae;o__Caldilineales;f__Caldilineaceae;g__uncultured | 0.09 | 0.12 | 0.00 | 0.00 | 0.00 | 0.00 | 0.00 | 0.00 | 0.00 |
| Chloroflexi;c__Anaerolineae;o__RBG-13-54-9;f__RBG-13-54-9;g__RBG-13-54-9 | 0.36 | 0.00 | 0.00 | 0.00 | 0.00 | 0.00 | 0.00 | 0.00 | 0.00 |
| Chloroflexi;c__Anaerolineae;o__SBR1031;f__A4b;g__A4b | 0.11 | 0.01 | 0.00 | 0.05 | 0.00 | 0.00 | 0.14 | 0.00 | 0.01 |
| Chloroflexi;c__Anaerolineae;o__SBR1031;f__SBR1031;g__SBR1031 | 0.15 | 0.00 | 0.00 | 0.00 | 0.00 | 0.00 | 0.00 | 0.00 | 0.00 |
| Chloroflexi;c__Chloroflexia;o__Chloroflexales;f__Roseiflexaceae;g__uncultured | 0.12 | 0.00 | 0.00 | 0.00 | 0.00 | 0.00 | 0.00 | 0.00 | 0.00 |
| Chloroflexi;c__Chloroflexia;o__Thermomicrobiales;f__JG30-KF-CM45;g__JG30-KF-CM45 | 0.63 | 0.00 | 0.00 | 0.00 | 0.00 | 0.08 | 0.00 | 0.00 | 0.00 |
| Chloroflexi;c__Dehalococcoidia;o__S085;f__S085;g__S085 | 0.15 | 0.00 | 0.00 | 0.00 | 0.00 | 0.00 | 0.00 | 0.00 | 0.00 |
| Chloroflexi;c__Gitt-GS-136;o__Gitt-GS-136;f__Gitt-GS-136;g__Gitt-GS-136 | 0.12 | 0.00 | 0.00 | 0.00 | 0.00 | 0.00 | 0.00 | 0.00 | 0.00 |
| Chloroflexi;c__JG30-KF-CM66;o__JG30-KF-CM66;f__JG30-KF-CM66;g__JG30-KF-CM66 | 0.07 | 0.02 | 0.00 | 0.00 | 0.00 | 0.03 | 0.49 | 0.00 | 0.00 |
| Chloroflexi;c__KD4-96;o__KD4-96;f__KD4-96;g__KD4-96 | 3.04 | 0.01 | 0.04 | 0.00 | 0.00 | 0.00 | 0.00 | 0.03 | 0.00 |
| Chloroflexi;c__Ktedonobacteria;o__C0119;f__C0119;g__C0119 | 0.25 | 0.00 | 0.00 | 0.00 | 0.00 | 0.00 | 0.00 | 0.00 | 0.00 |
| Chloroflexi;c__Ktedonobacteria;o__Ktedonobacterales;f__JG30-KF-AS9;g__JG30-KF-AS9 | 0.28 | 0.00 | 0.00 | 0.00 | 0.00 | 0.00 | 0.00 | 0.00 | 0.00 |
| Chloroflexi;c__Ktedonobacteria;o__Ktedonobacterales;f__Ktedonobacteraceae | 0.10 | 0.00 | 0.00 | 0.00 | 0.00 | 0.00 | 0.00 | 0.00 | 0.00 |
| Chloroflexi;c__Ktedonobacteria;o__Ktedonobacterales;f__Ktedonobacteraceae;g__uncultured | 0.10 | 0.00 | 0.00 | 0.00 | 0.00 | 0.00 | 0.00 | 0.00 | 0.00 |
| Chloroflexi;c__TK10;o__TK10;f__TK10;g__TK10 | 0.85 | 0.00 | 0.00 | 0.00 | 0.00 | 0.09 | 0.00 | 0.00 | 0.00 |
| Desulfobacterota;c__Desulfuromonadia;o__Geobacterales;f__Geobacteraceae | 0.14 | 0.00 | 0.00 | 0.00 | 0.00 | 0.00 | 0.00 | 0.00 | 0.00 |
| Desulfobacterota;c__uncultured;o__uncultured;f__uncultured;g__uncultured | 0.21 | 0.00 | 0.00 | 0.00 | 0.00 | 0.00 | 0.00 | 0.00 | 0.00 |
| Firmicutes;c__Bacilli;o__Bacillales;f__Bacillaceae;g__Bacillus | 0.74 | 0.00 | 0.00 | 0.00 | 0.00 | 0.00 | 0.10 | 0.00 | 0.00 |
| Firmicutes;c__Bacilli;o__Lactobacillales;f__Lactobacillaceae;g__Lactobacillus | 0.00 | 0.00 | 0.00 | 0.00 | 0.10 | 0.00 | 0.00 | 0.00 | 0.00 |
| Firmicutes;c__Bacilli;o__Paenibacillales;f__Paenibacillaceae;g__Paenibacillus | 0.07 | 0.00 | 0.00 | 0.00 | 0.00 | 0.00 | 0.00 | 0.00 | 0.00 |
| Firmicutes;c__Bacilli;o__Thermoactinomycetales;f__Thermoactinomycetaceae;g__Planifilum | 0.00 | 0.00 | 0.00 | 0.00 | 0.19 | 0.00 | 0.00 | 0.00 | 0.00 |
| Firmicutes;c__Clostridia;o__Clostridiales;f__Clostridiaceae | 0.10 | 0.00 | 0.00 | 0.00 | 0.00 | 0.00 | 0.00 | 0.00 | 0.00 |
| Gemmatimonadota;c__Gemmatimonadetes;o__Gemmatimonadales;f__Gemmatimonadaceae | 0.14 | 0.00 | 0.00 | 0.00 | 0.00 | 0.00 | 0.00 | 0.00 | 0.00 |
| Gemmatimonadota;c__Gemmatimonadetes;o__Gemmatimonadales;f__Gemmatimonadaceae;g__Gemmatimonas | 0.64 | 0.01 | 0.02 | 0.00 | 0.00 | 0.00 | 0.00 | 0.00 | 0.00 |
| Gemmatimonadota;c__Gemmatimonadetes;o__Gemmatimonadales;f__Gemmatimonadaceae;g__uncultured | 0.49 | 0.02 | 0.00 | 0.00 | 0.00 | 0.00 | 0.00 | 0.00 | 0.00 |
| Latescibacterota;c__Latescibacterota;o__Latescibacterota;f__Latescibacterota;g__Latescibacterota | 0.96 | 0.00 | 0.00 | 0.00 | 0.00 | 0.00 | 0.00 | 0.00 | 0.00 |
| Methylomirabilota;c__Methylomirabilia;o__Rokubacteriales;f__Rokubacteriales;g__Rokubacteriales | 0.16 | 0.00 | 0.00 | 0.00 | 0.00 | 0.00 | 0.00 | 0.00 | 0.00 |
| Methylomirabilota;c__Methylomirabilia;o__Rokubacteriales;f__WX65;g__WX65 | 0.33 | 0.00 | 0.00 | 0.00 | 0.00 | 0.00 | 0.00 | 0.00 | 0.00 |
| Myxococcota;c__Myxococcia;o__Myxococcales;f__Anaeromyxobacteraceae;g__Anaeromyxobacter | 0.17 | 0.00 | 0.00 | 0.00 | 0.00 | 0.00 | 0.00 | 0.00 | 0.00 |
| Myxococcota;c__Myxococcia;o__Myxococcales;f__Myxococcaceae | 0.14 | 0.00 | 0.00 | 0.00 | 0.00 | 0.00 | 0.00 | 0.00 | 0.00 |
| Myxococcota;c__Myxococcia;o__Myxococcales;f__Myxococcaceae;g__P3OB-42 | 0.08 | 0.00 | 0.00 | 0.00 | 0.00 | 0.00 | 0.00 | 0.00 | 0.00 |
| Myxococcota;c__Myxococcia;o__Myxococcales;f__Myxococcaceae;g__uncultured | 0.18 | 0.00 | 0.00 | 0.00 | 0.00 | 0.00 | 0.00 | 0.00 | 0.00 |
| Myxococcota;c__Polyangia;o__Haliangiales;f__Haliangiaceae;g__Haliangium | 0.84 | 0.00 | 0.00 | 0.00 | 0.00 | 0.00 | 0.00 | 0.00 | 0.00 |
| Myxococcota;c__Polyangia;o__Polyangiales;f__BIrii41;g__BIrii41 | 0.13 | 0.00 | 0.00 | 0.00 | 0.00 | 0.00 | 0.00 | 0.00 | 0.00 |
| Myxococcota;c__Polyangia;o__Polyangiales;f__Phaselicystidaceae;g__Phaselicystis | 0.22 | 0.00 | 0.00 | 0.00 | 0.00 | 0.00 | 0.00 | 0.00 | 0.00 |
| Myxococcota;c__Polyangia;o__Polyangiales;f__Polyangiaceae;g__Pajaroellobacter | 0.16 | 0.00 | 0.00 | 0.00 | 0.00 | 0.00 | 0.00 | 0.00 | 0.00 |
| Nitrospirota;c__Nitrospiria;o__Nitrospirales;f__Nitrospiraceae;g__Nitrospira | 0.36 | 0.00 | 0.00 | 0.00 | 0.00 | 0.00 | 0.00 | 0.00 | 0.00 |
| Patescibacteria;c__Saccharimonadia;o__Saccharimonadales;f__Saccharimonadales;g__Saccharimonadales | 0.08 | 0.00 | 0.05 | 0.00 | 0.00 | 0.00 | 0.01 | 0.00 | 0.00 |
| Planctomycetota;c__Planctomycetes;o__Gemmatales;f__Gemmataceae | 0.17 | 0.00 | 0.00 | 0.00 | 0.00 | 0.00 | 0.00 | 0.00 | 0.00 |
| Planctomycetota;c__Planctomycetes;o__Gemmatales;f__Gemmataceae;g__Fimbriiglobus | 0.06 | 0.00 | 0.00 | 0.00 | 0.00 | 0.00 | 0.00 | 0.00 | 0.00 |
| Planctomycetota;c__Planctomycetes;o__Gemmatales;f__Gemmataceae;g__Gemmata | 0.29 | 0.00 | 0.00 | 0.00 | 0.00 | 0.00 | 0.00 | 0.00 | 0.00 |
| Planctomycetota;c__Planctomycetes;o__Gemmatales;f__Gemmataceae;g__Telmatocola | 0.09 | 0.03 | 0.00 | 0.00 | 0.00 | 0.00 | 0.00 | 0.00 | 0.00 |
| Planctomycetota;c__Planctomycetes;o__Gemmatales;f__Gemmataceae;g__uncultured | 0.86 | 0.01 | 0.00 | 0.08 | 0.00 | 0.05 | 0.00 | 0.00 | 0.00 |
| Planctomycetota;c__Planctomycetes;o__Isosphaerales;f__Isosphaeraceae | 0.37 | 0.00 | 0.00 | 0.00 | 0.00 | 0.00 | 0.00 | 0.00 | 0.00 |
| Planctomycetota;c__Planctomycetes;o__Pirellulales;f__Pirellulaceae | 0.08 | 0.00 | 0.00 | 0.02 | 0.00 | 0.00 | 0.00 | 0.00 | 0.00 |
| Planctomycetota;c__Planctomycetes;o__Pirellulales;f__Pirellulaceae;g__Pir4_lineage | 0.10 | 0.00 | 0.00 | 0.00 | 0.00 | 0.00 | 0.00 | 0.00 | 0.00 |
| Planctomycetota;c__Planctomycetes;o__Pirellulales;f__Pirellulaceae;g__Pirellula | 0.17 | 0.01 | 0.00 | 0.00 | 0.00 | 0.07 | 0.00 | 0.00 | 0.00 |
| Planctomycetota;c__Planctomycetes;o__Planctomycetales;f__Schlesneriaceae;g__Schlesneria | 0.00 | 0.00 | 0.00 | 0.00 | 0.00 | 0.00 | 0.00 | 0.00 | 0.00 |
| Planctomycetota;c__Planctomycetes;o__Planctomycetales;f__uncultured;g__uncultured | 0.18 | 0.00 | 0.02 | 0.00 | 0.00 | 0.00 | 0.00 | 0.00 | 0.00 |
| Proteobacteria;c__Alphaproteobacteria | 0.51 | 0.00 | 0.10 | 0.12 | 0.00 | 0.00 | 0.00 | 0.00 | 0.00 |
| Proteobacteria;c__Alphaproteobacteria;o__Acetobacterales;f__Acetobacteraceae | 0.15 | 0.07 | 0.10 | 0.08 | 0.00 | 0.00 | 0.02 | 0.06 | 0.00 |
| Proteobacteria;c__Alphaproteobacteria;o__Azospirillales;f__Inquilinaceae;g__Inquilinus | 0.00 | 0.02 | 0.00 | 0.00 | 0.00 | 0.12 | 0.02 | 0.00 | 0.00 |
| Proteobacteria;c__Alphaproteobacteria;o__Azospirillales;f__uncultured;g__uncultured | 0.13 | 0.00 | 0.00 | 0.00 | 0.00 | 0.00 | 0.00 | 0.00 | 0.00 |
| Proteobacteria;c__Alphaproteobacteria;o__Caulobacterales;f__Caulobacteraceae | 0.02 | 0.99 | 1.38 | 0.20 | 0.10 | 0.37 | 0.00 | 0.39 | 0.05 |
| Proteobacteria;c__Alphaproteobacteria;o__Caulobacterales;f__Caulobacteraceae;g__Phenylobacterium | 0.10 | 0.34 | 0.12 | 0.11 | 0.17 | 0.00 | 0.00 | 0.00 | 0.00 |
| Proteobacteria;c__Alphaproteobacteria;o__Caulobacterales;f__Caulobacteraceae;g__uncultured | 0.02 | 0.08 | 0.03 | 0.00 | 0.00 | 0.00 | 0.00 | 0.00 | 0.00 |
| Proteobacteria;c__Alphaproteobacteria;o__Defluviicoccales;f__Defluviicoccaceae;g__Defluviicoccus | 0.13 | 0.00 | 0.00 | 0.00 | 0.00 | 0.00 | 0.00 | 0.00 | 0.00 |
| Proteobacteria;c__Alphaproteobacteria;o__Dongiales;f__Dongiaceae;g__Dongia | 0.18 | 0.02 | 0.00 | 0.00 | 0.00 | 0.00 | 0.00 | 0.00 | 0.00 |
| Proteobacteria;c__Alphaproteobacteria;o__Elsterales;f__uncultured;g__uncultured | 0.03 | 0.01 | 0.06 | 0.02 | 0.00 | 0.00 | 0.02 | 0.00 | 0.00 |
| Proteobacteria;c__Alphaproteobacteria;o__Reyranellales;f__Reyranellaceae;g__Reyranella | 0.73 | 0.59 | 0.58 | 0.28 | 0.03 | 0.79 | 0.13 | 0.00 | 0.04 |
| Proteobacteria;c__Alphaproteobacteria;o__Rhizobiales | 0.21 | 0.22 | 0.00 | 0.11 | 0.00 | 0.21 | 0.05 | 0.04 | 0.00 |
| Proteobacteria;c__Alphaproteobacteria;o__Rhizobiales;f__Beijerinckiaceae | 0.17 | 0.02 | 0.02 | 0.04 | 0.00 | 0.00 | 0.06 | 0.00 | 0.00 |
| Proteobacteria;c__Alphaproteobacteria;o__Rhizobiales;f__Beijerinckiaceae;g__Methylobacterium-Methylorubrum | 0.01 | 0.22 | 0.03 | 0.00 | 0.07 | 0.16 | 0.00 | 0.00 | 0.00 |
| Proteobacteria;c__Alphaproteobacteria;o__Rhizobiales;f__Devosiaceae | 0.00 | 0.09 | 0.06 | 0.07 | 0.05 | 0.37 | 0.24 | 0.01 | 0.08 |
| Proteobacteria;c__Alphaproteobacteria;o__Rhizobiales;f__Devosiaceae;g__Devosia | 0.02 | 0.09 | 0.10 | 0.00 | 0.00 | 0.00 | 0.00 | 0.06 | 0.03 |
| Proteobacteria;c__Alphaproteobacteria;o__Rhizobiales;f__Hyphomicrobiaceae | 0.31 | 0.00 | 0.00 | 0.00 | 0.00 | 0.00 | 0.00 | 0.00 | 0.00 |
| Proteobacteria;c__Alphaproteobacteria;o__Rhizobiales;f__Hyphomicrobiaceae;g__Pedomicrobium | 0.42 | 0.00 | 0.00 | 0.00 | 0.00 | 0.00 | 0.00 | 0.00 | 0.00 |
| Proteobacteria;c__Alphaproteobacteria;o__Rhizobiales;f__Labraceae;g__Labrys | 0.00 | 0.00 | 0.00 | 0.07 | 0.00 | 0.00 | 0.01 | 0.00 | 0.02 |
| Proteobacteria;c__Alphaproteobacteria;o__Rhizobiales;f__Methyloligellaceae;g__Methyloceanibacter | 0.71 | 0.00 | 0.00 | 0.00 | 0.00 | 0.00 | 0.00 | 0.00 | 0.00 |
| Proteobacteria;c__Alphaproteobacteria;o__Rhizobiales;f__Methyloligellaceae;g__uncultured | 1.64 | 0.00 | 0.00 | 0.00 | 0.00 | 0.00 | 0.00 | 0.00 | 0.00 |
| Proteobacteria;c__Alphaproteobacteria;o__Rhizobiales;f__Rhizobiaceae | 0.34 | 0.38 | 0.70 | 0.42 | 0.00 | 1.36 | 2.26 | 0.56 | 0.07 |
| Proteobacteria;c__Alphaproteobacteria;o__Rhizobiales;f__Rhizobiaceae;g__Allorhizobium-Neorhizobium-Pararhizobium-Rhiz | 0.08 | 1.20 | 1.61 | 3.44 | 0.00 | 7.18 | 6.65 | 0.76 | 0.54 |
| Proteobacteria;c__Alphaproteobacteria;o__Rhizobiales;f__Rhizobiales_Incertae_Sedis;g__Nordella | 0.12 | 0.00 | 0.00 | 0.00 | 0.00 | 0.00 | 0.00 | 0.00 | 0.00 |
| Proteobacteria;c__Alphaproteobacteria;o__Rhizobiales;f__Rhodomicrobiaceae;g__Rhodomicrobium | 0.15 | 0.00 | 0.00 | 0.00 | 0.00 | 0.00 | 0.00 | 0.00 | 0.00 |
| Proteobacteria;c__Alphaproteobacteria;o__Rhizobiales;f__Xanthobacteraceae | 1.05 | 2.12 | 1.96 | 2.78 | 5.24 | 5.04 | 1.96 | 2.13 | 1.43 |
| Proteobacteria;c__Alphaproteobacteria;o__Rhizobiales;f__Xanthobacteraceae | 9.39 | 2.51 | 2.31 | 3.24 | 5.62 | 5.64 | 2.31 | 2.28 | 1.66 |
| Proteobacteria;c__Alphaproteobacteria;o__Rhizobiales;f__Xanthobacteraceae;g__Bradyrhizobium | 0.90 | 0.37 | 0.26 | 0.40 | 0.38 | 0.44 | 0.29 | 0.15 | 0.21 |
| Proteobacteria;c__Alphaproteobacteria;o__Rhizobiales;f__Xanthobacteraceae;g__Pseudolabrys | 0.03 | 0.00 | 0.00 | 0.03 | 0.00 | 0.00 | 0.06 | 0.00 | 0.00 |
| Proteobacteria;c__Alphaproteobacteria;o__Rhizobiales;f__Xanthobacteraceae;g__Rhodoplanes | 0.33 | 0.02 | 0.00 | 0.00 | 0.00 | 0.00 | 0.00 | 0.00 | 0.00 |
| Proteobacteria;c__Alphaproteobacteria;o__Rhizobiales;f__Xanthobacteraceae;g__uncultured | 7.08 | 0.00 | 0.10 | 0.03 | 0.00 | 0.16 | 0.00 | 0.00 | 0.02 |
| Proteobacteria;c__Alphaproteobacteria;o__Rhodobacterales;f__Rhodobacteraceae;g__Rhodobacter | 0.00 | 0.05 | 0.00 | 0.09 | 0.00 | 0.00 | 0.01 | 0.00 | 0.06 |
| Proteobacteria;c__Alphaproteobacteria;o__Sphingomonadales;f__Sphingomonadaceae | 0.40 | 40.27 | 35.67 | 49.32 | 90.78 | 28.67 | 15.85 | 6.55 | 18.00 |
| Proteobacteria;c__Alphaproteobacteria;o__Sphingomonadales;f__Sphingomonadaceae;g__Novosphingobium | 0.00 | 0.74 | 0.62 | 0.17 | 0.07 | 1.22 | 0.01 | 0.06 | 0.57 |
| Proteobacteria;c__Alphaproteobacteria;o__Sphingomonadales;f__Sphingomonadaceae;g__Sphingobium | 0.00 | 29.03 | 15.66 | 16.69 | 0.34 | 13.03 | 4.23 | 3.12 | 1.40 |
| Proteobacteria;c__Alphaproteobacteria;o__Sphingomonadales;f__Sphingomonadaceae;g__Sphingomonas | 0.06 | 2.91 | 1.48 | 1.48 | 0.05 | 0.35 | 0.17 | 0.37 | 2.02 |
| Proteobacteria;c__Alphaproteobacteria;o__Tistrellales;f__Geminicoccaceae;g__Candidatus_Alysiosphaera | 0.11 | 0.00 | 0.00 | 0.00 | 0.00 | 0.00 | 0.00 | 0.00 | 0.00 |
| Proteobacteria;c__Alphaproteobacteria;o__uncultured;f__uncultured;g__uncultured | 0.15 | 0.00 | 0.03 | 0.00 | 0.00 | 0.00 | 0.00 | 0.00 | 0.00 |
| Proteobacteria;c__Gammaproteobacteria | 0.33 | 0.03 | 0.02 | 0.00 | 0.00 | 0.00 | 0.00 | 0.00 | 0.00 |
| Proteobacteria;c__Gammaproteobacteria;o__Burkholderiales | 0.24 | 0.00 | 0.00 | 0.00 | 0.00 | 0.00 | 0.00 | 0.00 | 0.00 |
| Proteobacteria;c__Gammaproteobacteria;o__Burkholderiales;f__A21b;g__A21b | 0.22 | 0.00 | 0.00 | 0.00 | 0.00 | 0.00 | 0.00 | 0.00 | 0.00 |
| Proteobacteria;c__Gammaproteobacteria;o__Burkholderiales;f__Burkholderiaceae | 0.06 | 0.82 | 1.27 | 0.24 | 0.13 | 2.90 | 0.61 | 0.51 | 0.22 |
| Proteobacteria;c__Gammaproteobacteria;o__Burkholderiales;f__Burkholderiaceae;g__Burkholderia-Caballeronia-Paraburkho | 0.04 | 0.25 | 0.17 | 0.11 | 0.00 | 2.00 | 0.26 | 0.13 | 0.06 |
| Proteobacteria;c__Gammaproteobacteria;o__Burkholderiales;f__Burkholderiaceae;g__Cupriavidus | 0.02 | 0.21 | 0.72 | 0.07 | 0.06 | 0.16 | 0.26 | 0.23 | 0.16 |
| Proteobacteria;c__Gammaproteobacteria;o__Burkholderiales;f__Burkholderiaceae;g__Ralstonia | 0.00 | 0.36 | 0.38 | 0.06 | 0.08 | 0.73 | 0.09 | 0.16 | 0.00 |
| Proteobacteria;c__Gammaproteobacteria;o__Burkholderiales;f__Comamonadaceae | 0.14 | 0.52 | 0.64 | 0.14 | 0.05 | 0.05 | 0.06 | 0.10 | 0.01 |
| Proteobacteria;c__Gammaproteobacteria;o__Burkholderiales;f__Comamonadaceae;g__Ramlibacter | 0.05 | 0.03 | 0.03 | 0.00 | 0.00 | 0.09 | 0.00 | 0.05 | 0.00 |
| Proteobacteria;c__Gammaproteobacteria;o__Burkholderiales;f__Comamonadaceae;g__Rhodoferax | 0.00 | 0.53 | 0.17 | 0.39 | 0.00 | 0.07 | 0.00 | 0.55 | 0.06 |
| Proteobacteria;c__Gammaproteobacteria;o__Burkholderiales;f__Comamonadaceae;g__Rubrivivax | 0.04 | 0.05 | 0.03 | 0.02 | 0.00 | 0.00 | 0.00 | 0.00 | 0.00 |
| Proteobacteria;c__Gammaproteobacteria;o__Burkholderiales;f__Comamonadaceae;g__Variovorax | 0.02 | 1.54 | 0.80 | 0.02 | 0.00 | 0.79 | 0.13 | 0.55 | 0.21 |
| Proteobacteria;c__Gammaproteobacteria;o__Burkholderiales;f__Nitrosomonadaceae;g__Ellin6067 | 0.40 | 0.00 | 0.05 | 0.00 | 0.00 | 0.00 | 0.00 | 0.00 | 0.00 |
| Proteobacteria;c__Gammaproteobacteria;o__Burkholderiales;f__Nitrosomonadaceae;g__mle1-7 | 0.21 | 0.00 | 0.00 | 0.00 | 0.00 | 0.00 | 0.00 | 0.00 | 0.00 |
| Proteobacteria;c__Gammaproteobacteria;o__Burkholderiales;f__Nitrosomonadaceae;g__MND1 | 0.40 | 0.00 | 0.02 | 0.00 | 0.00 | 0.00 | 0.00 | 0.00 | 0.00 |
| Proteobacteria;c__Gammaproteobacteria;o__Burkholderiales;f__SC-I-84;g__SC-I-84 | 0.92 | 0.00 | 0.00 | 0.00 | 0.00 | 0.00 | 0.00 | 0.00 | 0.00 |
| Proteobacteria;c__Gammaproteobacteria;o__CCD24;f__CCD24;g__CCD24 | 0.14 | 0.00 | 0.00 | 0.00 | 0.00 | 0.00 | 0.00 | 0.00 | 0.00 |
| Proteobacteria;c__Gammaproteobacteria;o__Diplorickettsiales;f__Diplorickettsiaceae;g__Aquicella | 0.09 | 0.00 | 0.00 | 0.00 | 0.01 | 0.00 | 0.00 | 0.00 | 0.00 |
| Proteobacteria;c__Gammaproteobacteria;o__Diplorickettsiales;f__Diplorickettsiaceae;g__uncultured | 0.06 | 0.02 | 0.00 | 0.03 | 0.00 | 0.00 | 0.00 | 0.00 | 0.00 |
| Proteobacteria;c__Gammaproteobacteria;o__Enterobacterales | 0.00 | 0.05 | 0.07 | 0.83 | 0.00 | 0.00 | 0.71 | 0.04 | 0.06 |
| Proteobacteria;c__Gammaproteobacteria;o__Enterobacterales;f__Enterobacteriaceae | 0.00 | 0.23 | 0.12 | 0.20 | 0.07 | 0.33 | 3.08 | 0.00 | 0.28 |
| Proteobacteria;c__Gammaproteobacteria;o__Enterobacterales;f__Enterobacteriaceae | 0.00 | 0.26 | 0.12 | 1.43 | 0.07 | 0.73 | 4.50 | 0.05 | 0.32 |
| Proteobacteria;c__Gammaproteobacteria;o__Enterobacterales;f__Enterobacteriaceae;g__Enterobacter | 0.00 | 0.00 | 0.00 | 0.13 | 0.00 | 0.00 | 0.37 | 0.05 | 0.00 |
| Proteobacteria;c__Gammaproteobacteria;o__Enterobacterales;f__Erwiniaceae | 0.00 | 0.03 | 0.00 | 0.97 | 0.00 | 0.40 | 1.01 | 0.00 | 0.04 |
| Proteobacteria;c__Gammaproteobacteria;o__Enterobacterales;f__Erwiniaceae;g__Erwinia | 0.00 | 0.00 | 0.00 | 0.14 | 0.00 | 0.00 | 0.03 | 0.00 | 0.00 |
| Proteobacteria;c__Gammaproteobacteria;o__Enterobacterales;f__Erwiniaceae;g__Pantoea | 0.00 | 0.06 | 0.26 | 3.10 | 0.00 | 0.24 | 0.26 | 0.00 | 0.00 |
| Proteobacteria;c__Gammaproteobacteria;o__Enterobacterales;f__Yersiniaceae;g__Rahnella1 | 0.00 | 0.00 | 0.01 | 1.20 | 0.00 | 0.98 | 0.90 | 0.14 | 0.32 |
| Proteobacteria;c__Gammaproteobacteria;o__Enterobacterales;f__Yersiniaceae;g__Serratia | 0.00 | 0.00 | 0.01 | 0.74 | 0.00 | 0.00 | 0.16 | 0.00 | 0.00 |
| Proteobacteria;c__Gammaproteobacteria;o__Gammaproteobacteria_Incertae_Sedis;f__Unknown_Family;g__Acidibacter | 0.21 | 0.00 | 0.00 | 0.00 | 0.00 | 0.00 | 0.00 | 0.00 | 0.00 |
| Proteobacteria;c__Gammaproteobacteria;o__Pseudomonadales;f__Moraxellaceae;g__Acinetobacter | 0.00 | 0.05 | 0.06 | 0.03 | 0.00 | 0.05 | 0.00 | 0.58 | 0.03 |
| Proteobacteria;c__Gammaproteobacteria;o__Pseudomonadales;f__Pseudomonadaceae;g__Pseudomonas | 0.06 | 2.97 | 5.82 | 2.25 | 0.02 | 3.27 | 24.48 | 69.56 | 60.16 |
| Proteobacteria;c__Gammaproteobacteria;o__Steroidobacterales;f__Steroidobacteraceae;g__Steroidobacter | 0.18 | 0.00 | 0.00 | 0.00 | 0.00 | 0.00 | 0.00 | 0.00 | 0.00 |
| Proteobacteria;c__Gammaproteobacteria;o__Steroidobacterales;f__Steroidobacteraceae;g__uncultured | 0.10 | 0.00 | 0.00 | 0.00 | 0.00 | 0.00 | 0.00 | 0.00 | 0.00 |
| Proteobacteria;c__Gammaproteobacteria;o__Xanthomonadales;f__Xanthomonadaceae;g__Stenotrophomonas | 0.00 | 0.10 | 0.04 | 0.02 | 0.00 | 0.04 | 0.00 | 0.34 | 0.03 |
| Verrucomicrobiota;c__Verrucomicrobiae;o__Chthoniobacterales | 1.85 | 0.00 | 0.00 | 0.00 | 0.00 | 0.00 | 0.00 | 0.00 | 0.02 |
| Verrucomicrobiota;c__Verrucomicrobiae;o__Chthoniobacterales;f__Chthoniobacteraceae;g__Candidatus_Udaeobacter | 9.39 | 0.00 | 0.00 | 0.00 | 0.00 | 0.00 | 0.00 | 0.00 | 0.00 |
| Verrucomicrobiota;c__Verrucomicrobiae;o__Chthoniobacterales;f__Chthoniobacteraceae;g__Chthoniobacter | 0.32 | 0.20 | 0.02 | 0.08 | 0.00 | 0.05 | 0.04 | 0.08 | 0.07 |
| Verrucomicrobiota;c__Verrucomicrobiae;o__Chthoniobacterales;f__Terrimicrobiaceae;g__Terrimicrobium | 0.24 | 0.00 | 0.00 | 0.00 | 0.00 | 0.15 | 0.00 | 0.00 | 0.01 |
| Verrucomicrobiota;c__Verrucomicrobiae;o__Chthoniobacterales;f__Xiphinematobacteraceae;g__Candidatus_Xiphinematoba | 0.64 | 0.04 | 0.00 | 0.00 | 0.00 | 0.05 | 0.00 | 0.04 | 0.00 |
| Verrucomicrobiota;c__Verrucomicrobiae;o__Opitutales;f__Opitutaceae | 0.09 | 0.06 | 0.00 | 0.01 | 0.00 | 0.00 | 0.00 | 0.00 | 0.00 |
| Verrucomicrobiota;c__Verrucomicrobiae;o__Pedosphaerales;f__Pedosphaeraceae | 0.30 | 0.00 | 0.00 | 0.00 | 0.00 | 0.00 | 0.00 | 0.00 | 0.00 |
| Verrucomicrobiota;c__Verrucomicrobiae;o__Pedosphaerales;f__Pedosphaeraceae;g__ADurb.Bin063-1 | 0.63 | 0.00 | 0.00 | 0.00 | 0.00 | 0.00 | 0.00 | 0.00 | 0.00 |
| Verrucomicrobiota;c__Verrucomicrobiae;o__Pedosphaerales;f__Pedosphaeraceae;g__Pedosphaeraceae | 0.58 | 0.00 | 0.00 | 0.00 | 0.00 | 0.00 | 0.00 | 0.00 | 0.00 |
| Verrucomicrobiota;c__Verrucomicrobiae;o__Pedosphaerales;f__Pedosphaeraceae;g__uncultured | 0.26 | 0.00 | 0.00 | 0.00 | 0.00 | 0.00 | 0.00 | 0.00 | 0.00 |
| Verrucomicrobiota;c__Verrucomicrobiae;o__Verrucomicrobiales;f__Rubritaleaceae;g__Luteolibacter | 0.05 | 0.00 | 0.00 | 0.00 | 0.00 | 0.00 | 0.00 | 0.00 | 0.15 |
| Verrucomicrobiota;c__Verrucomicrobiae;o__Verrucomicrobiales;f__Verrucomicrobiaceae | 0.01 | 0.08 | 0.00 | 0.00 | 0.00 | 0.00 | 0.00 | 0.00 | 0.14 |
| Verrucomicrobiota;c__Verrucomicrobiae;o__Verrucomicrobiales;f__Verrucomicrobiaceae;g__Roseimicrobium | 0.12 | 0.05 | 0.00 | 0.12 | 0.00 | 0.08 | 0.00 | 0.01 | 0.01 |
| Verrucomicrobiota;c__Verrucomicrobiae;o__Verrucomicrobiales;f__Verrucomicrobiaceae;g__uncultured | 0.03 | 0.03 | 0.00 | 0.00 | 0.00 | 0.00 | 0.00 | 0.00 | 0.14 |
| d__Archaea;Crenarchaeota;c__Nitrososphaeria;o__Nitrososphaerales;f__Nitrososphaeraceae;g__Nitrososphaeraceae | 1.16 | 0.00 | 0.00 | 0.00 | 0.00 | 0.00 | 0.00 | 0.00 | 0.00 |

**Relative taxon abundance in Arctic core samples and in permafrost populations sorted follwing BONCAT incubations.**

D1, active layer; D3, permafrost; G, sorted based on A488 click label; N, incubation with exogenous nutrients

| Taxon | D1 | D3 | D3_6w G | D3_6wN G |
|---|---|---|---|---|
| Acidobacteriota;c__Holophagae;o__Subgroup_7;f__Subgroup_7;g__Subgroup_7 | 1.16 | 0.00 | 0.00 | 0.00 |
| Acidobacteriota;c__Vicinamibacteria;o__Vicinamibacterales;f__uncultured;g__uncultured | 7.48 | 0.00 | 0.00 | 0.00 |
| Acidobacteriota;c__Vicinamibacteria;o__Vicinamibacterales;f__Vicinamibacteraceae;g__Vicinamibacteraceae | 0.93 | 0.00 | 0.00 | 0.00 |
| Actinobacteriota;c__Acidimicrobiia;o__IMCC26256;f__IMCC26256;g__IMCC26256 | 0.87 | 0.77 | 0.00 | 0.00 |
| Actinobacteriota;c__Acidimicrobiia;o__Microtrichales;f__Ilumatobacteraceae;g__CL500-29_marine_group | 3.13 | 0.00 | 0.00 | 0.00 |
| Actinobacteriota;c__Actinobacteria;o__0319-7L14;f__0319-7L14;g__0319-7L14 | 0.00 | 6.12 | 0.00 | 0.00 |
| Actinobacteriota;c__Actinobacteria;o__Corynebacteriales;f__Corynebacteriaceae;g__Corynebacterium | 0.23 | 1.01 | 0.00 | 0.00 |
| Actinobacteriota;c__Actinobacteria;o__Corynebacteriales;f__Corynebacteriaceae;g__Lawsonella | 0.00 | 2.35 | 0.00 | 0.03 |
| Actinobacteriota;c__Actinobacteria;o__Corynebacteriales;f__Nocardiaceae;g__Rhodococcus | 0.44 | 1.84 | 59.13 | 31.60 |
| Actinobacteriota;c__Actinobacteria;o__Micrococcales;f__Microbacteriaceae;__ | 0.00 | 0.00 | 0.00 | 0.45 |
| Actinobacteriota;c__Actinobacteria;o__Micrococcales;f__Microbacteriaceae;g__Leifsonia | 0.00 | 0.00 | 0.00 | 0.00 |
| Actinobacteriota;c__Actinobacteria;o__Micrococcales;f__Micrococcaceae;__ | 1.30 | 0.00 | 0.00 | 6.31 |
| Actinobacteriota;c__Actinobacteria;o__Micrococcales;f__Micrococcaceae;g__Pseudarthrobacter | 4.46 | 0.00 | 0.00 | 0.00 |
| Actinobacteriota;c__Thermoleophilia;o__Gaiellales;f__Gaiellaceae;g__Gaiella | 1.81 | 0.00 | 0.00 | 0.00 |
| Actinobacteriota;c__Thermoleophilia;o__Gaiellales;f__uncultured;g__uncultured | 2.79 | 1.00 | 0.00 | 0.00 |
| Bacteroidota;c__Bacteroidia;o__Cytophagales;f__Hymenobacteraceae;g__Hymenobacter | 0.00 | 0.00 | 0.00 | 0.00 |
| Bacteroidota;c__Bacteroidia;o__Cytophagales;f__Spirosomaceae;g__Dyadobacter | 0.00 | 0.00 | 0.00 | 0.00 |
| Bacteroidota;c__Bacteroidia;o__Flavobacteriales;f__Flavobacteriaceae;g__Flavobacterium | 0.02 | 0.00 | 1.03 | 0.00 |
| Bacteroidota;c__Bacteroidia;o__Sphingobacteriales;f__Sphingobacteriaceae;g__Mucilaginibacter | 0.00 | 0.00 | 0.26 | 0.00 |
| Chloroflexi;c__Chloroflexia;o__Thermomicrobiales;f__JG30-KF-CM45;g__JG30-KF-CM45 | 0.07 | 0.00 | 0.00 | 0.00 |
| Chloroflexi;c__Gitt-GS-136;o__Gitt-GS-136;f__Gitt-GS-136;g__Gitt-GS-136 | 2.60 | 0.00 | 0.00 | 0.00 |
| Chloroflexi;c__KD4-96;o__KD4-96;f__KD4-96;g__KD4-96 | 4.90 | 0.00 | 0.00 | 0.00 |
| Chloroflexi;c__P2-11E;o__P2-11E;f__P2-11E;g__P2-11E | 0.86 | 0.59 | 0.00 | 0.00 |
| Firmicutes;c__Bacilli;o__Paenibacillales;f__Paenibacillaceae;g__Paenibacillus | 0.00 | 0.02 | 0.00 | 0.93 |
| Gemmatimonadota;c__Gemmatimonadetes;o__Gemmatimonadales;f__Gemmatimonadaceae;g__uncultured | 1.56 | 0.00 | 0.00 | 0.00 |
| Myxococcota;c__Polyangia;o__Nannocystales;f__Nannocystaceae;g__uncultured | 0.00 | 2.76 | 0.00 | 0.00 |
| Proteobacteria;c__Alphaproteobacteria;o__Rhizobiales;f__Rhizobiaceae;g__Aliihoeflea | 0.00 | 0.00 | 20.31 | 18.45 |
| Proteobacteria;c__Alphaproteobacteria;o__Rhizobiales;f__Rhizobiaceae;g__Aureimonas | 0.00 | 0.70 | 3.74 | 10.24 |
| Proteobacteria;c__Alphaproteobacteria;o__Rhizobiales;f__Rhizobiaceae;g__Phyllobacterium | 0.61 | 47.48 | 0.00 | 0.00 |
| Proteobacteria;c__Alphaproteobacteria;o__Rhizobiales;f__Xanthobacteraceae;g__Bradyrhizobium | 0.35 | 2.32 | 0.00 | 0.00 |
| Proteobacteria;c__Alphaproteobacteria;o__Rhizobiales;f__Xanthobacteraceae;g__uncultured | 11.89 | 0.00 | 0.00 | 0.00 |
| Proteobacteria;c__Alphaproteobacteria;o__Sphingomonadales;f__Sphingomonadaceae;g__Sphingomonas | 19.46 | 0.00 | 0.08 | 2.83 |
| Proteobacteria;c__Gammaproteobacteria;o__Burkholderiales;f__Comamonadaceae;g__Variovorax | 0.04 | 0.12 | 1.85 | 5.33 |
| Proteobacteria;c__Gammaproteobacteria;o__Burkholderiales;f__Oxalobacteraceae;g__Massilia | 0.00 | 1.33 | 7.43 | 0.00 |
| Proteobacteria;c__Gammaproteobacteria;o__Pseudomonadales;f__Moraxellaceae;g__Enhydrobacter | 0.00 | 3.60 | 0.00 | 0.00 |
| Proteobacteria;c__Gammaproteobacteria;o__Pseudomonadales;f__Moraxellaceae;g__Psychrobacter | 0.00 | 0.00 | 1.16 | 21.93 |
| Proteobacteria;c__Gammaproteobacteria;o__Pseudomonadales;f__Pseudomonadaceae;g__Pseudomonas | 0.79 | 9.47 | 2.97 | 0.08 |
| Verrucomicrobiota;c__Verrucomicrobiae;o__Chthoniobacterales;f__Chthoniobacteraceae;g__Candidatus_Udaeobacter | 1.42 | 0.00 | 0.00 | 0.00 |
| WPS-2;c__WPS-2;o__WPS-2;f__WPS-2;g__WPS-2 | 4.63 | 0.00 | 0.00 | 0.00 |

**Relative major genera/families abundance in oral sample and in populations sorted follwing BONCAT incubations.**

R, sorted based on red fluoresence (DNA stain). G, sorted based on A488 click label

| Description | Oral | MTGE R | MTGE G | MTGE High G | Dextrose R | Dextrose G | Lactate G | Lactate High G | NAG G |
|---|---|---|---|---|---|---|---|---|---|
| Actinobacteriota_Actinomyces | 2.98 | 0.05 | 0.13 | 0.05 | 0.38 | 0.24 | 0.18 | 0.03 | 0.34 |
| Actinobacteriota_Scardovia | 0.00 | 0.00 | 0.00 | 0.00 | 0.00 | 0.01 | 0.00 | 0.00 | 0.00 |
| Actinobacteriota_Corynebacterium | 0.41 | 0.00 | 0.00 | 0.00 | 0.03 | 0.02 | 0.01 | 0.01 | 0.00 |
| Actinobacteriota_Atopobiaceae | 0.01 | 0.46 | 0.83 | 0.57 | 5.79 | 8.78 | 6.46 | 4.60 | 5.94 |
| Bacteroidota_Paludibacteraceae | 0.00 | 0.15 | 0.04 | 0.33 | 0.55 | 0.77 | 0.32 | 2.71 | 0.23 |
| Bacteroidota_Porphyromonas | 1.07 | 0.10 | 0.61 | 0.13 | 0.96 | 1.14 | 0.62 | 0.24 | 0.71 |
| Bacteroidota_Alloprevotella | 6.09 | 0.48 | 1.15 | 1.69 | 0.58 | 0.69 | 0.49 | 0.28 | 0.85 |
| Bacteroidota_Prevotella | 19.97 | 2.56 | 4.08 | 5.92 | 9.50 | 10.61 | 7.57 | 15.43 | 9.57 |
| Bacteroidota_Capnocytophaga | 0.58 | 0.18 | 0.89 | 0.90 | 3.07 | 2.96 | 1.96 | 1.70 | 1.83 |
| Bacteroidota_Bergeyella | 0.00 | 0.29 | 0.01 | 0.51 | 0.76 | 1.44 | 0.52 | 5.63 | 0.28 |
| Campilobacterota_Campylobacter | 0.40 | 0.56 | 0.64 | 0.90 | 3.30 | 3.37 | 0.90 | 1.97 | 1.11 |
| Firmicutes_Gemella | 3.04 | 7.85 | 25.30 | 9.20 | 3.27 | 1.59 | 1.34 | 0.30 | 2.73 |
| Firmicutes_Granulicatella | 3.48 | 0.26 | 1.07 | 0.39 | 0.24 | 0.24 | 0.07 | 0.00 | 0.29 |
| Firmicutes_Streptococcus | 28.46 | 16.16 | 15.60 | 10.43 | 7.55 | 8.73 | 3.25 | 1.69 | 14.29 |
| Firmicutes_Lachnospiraceae | 1.88 | 6.30 | 0.74 | 3.66 | 1.77 | 3.86 | 0.98 | 5.89 | 2.35 |
| Firmicutes_Peptostreptococcus | 0.64 | 7.34 | 6.73 | 8.94 | 8.31 | 2.27 | 0.40 | 0.12 | 0.32 |
| Firmicutes_Selenomonas | 0.09 | 0.35 | 0.25 | 0.51 | 1.44 | 1.53 | 2.44 | 1.89 | 4.74 |
| Firmicutes_Veillonella | 7.59 | 17.45 | 22.40 | 7.94 | 16.28 | 6.87 | 56.48 | 11.84 | 28.66 |
| Fusobacteriota_Fusobacterium | 2.63 | 2.93 | 1.91 | 2.08 | 12.44 | 20.91 | 3.57 | 5.07 | 6.62 |
| Fusobacteriota_Leptotrichia | 1.29 | 24.74 | 0.43 | 20.48 | 3.51 | 3.82 | 1.03 | 9.46 | 2.70 |
| Patescibacteria_Saccharimonadaceae | 0.71 | 0.43 | 0.33 | 0.33 | 2.50 | 4.26 | 1.75 | 6.23 | 1.74 |
| Gammaproteobacteria_Neisseria | 2.57 | 0.00 | 0.02 | 0.00 | 0.43 | 0.28 | 0.26 | 0.13 | 0.22 |
| Gammaproteobacteria_Pasteurellaceae | 10.46 | 7.37 | 13.17 | 8.93 | 6.06 | 4.01 | 1.77 | 4.65 | 2.78 |
